# The NarX-NarL two-component system is a global regulator of biofilm formation, natural product biosynthesis, and host-associated survival in *Burkholderia pseudomallei*

**DOI:** 10.1101/2020.06.25.170712

**Authors:** Mihnea R. Mangalea, Bradley R. Borlee

**Author notes:** Address for correspondence: Brad Borlee.

## Abstract

In the environment, *Burkholderia pseudomallei* exists as a saprophyte inhabiting soils and surface waters where denitrification is important for anaerobic respiration. As an opportunistic pathogen, *B. pseudomallei* transitions from the environment to infect human and animal hosts where respiratory nitrate reduction enables replication in anoxic conditions. We have previously shown that *B. pseudomallei* responds to nitrate and nitrite in part by inhibiting biofilm formation and altering cyclic di-GMP signaling. Here, we describe the global transcriptomic response to nitrate and nitrite to characterize the nitrosative stress response relative to biofilm inhibition. To better understand the roles of nitrate-sensing in the biofilm inhibitory phenotype of *B. pseudomallei*, we created in-frame deletions of *narX* (Bp1026b_I1014) and *narL* (Bp1026b_I1013), which are adjacent components of the conserved nitrate-sensing two-component system. Through differential expression analysis of RNA-seq data, we observed that key components of the biofilm matrix are downregulated in response to nitrate and nitrite. In addition, several gene loci associated with the stringent response, central metabolism dysregulation, antibiotic tolerance, and pathogenicity determinants were significantly altered in their expression. Some of the most differentially expressed genes were nonribosomal peptide synthases (NRPS) and/or polyketide synthases (PKS) encoding the proteins for the biosynthesis of bactobolin, malleilactone, and syrbactin, in addition to an uncharacterized cryptic NRPS biosynthetic cluster. We also observed reduced expression of ribosomal structural and biogenesis loci, and gene clusters associated with translation and DNA replication, indicating modulation of growth rate and metabolism under nitrosative stress conditions. The differences in expression observed under nitrosative stress were reversed in *narX* and *narL* mutants, suggesting that nitrate sensing is an important checkpoint for regulating the diverse metabolic changes occurring in the biofilm inhibitory phenotype. Moreover, in a macrophage model of infection, *narX* and *narL* mutants were attenuated in intracellular replication, suggesting that nitrate sensing is important for host survival.

**Author Summary:** *Burkholderia pseudomallei* is a saprophytic bacterium inhabiting soils and surface waters throughout the tropics causing severe disease in humans and animals. Environmental signals such as the accumulation of inorganic ions mediates the biofilm forming capabilities and survival of *B. pseudomallei*. In particular, nitrate metabolism inhibits *B. pseudomallei* biofilm formation through complex regulatory cascades that relay environmental cues to intracellular second messengers that modulate bacterial physiology. Nitrates are common environmental contaminants derived from artificial fertilizers and byproducts of animal wastes that can be readily reduced by bacteria capable of denitrification. In *B. pseudomallei* 1026b, biofilm dynamics are in part regulated by a gene pathway involved in nitrate sensing, metabolism, and transport. This study investigated the role of a two-component nitrate sensing system, NarX-NarL, in regulating gene expression, biofilm formation, and cellular invasion. Global gene expression analyses in the wild type, as compared to Δ *narX* and Δ *narL* mutant strains with nitrate or nitrite implicate the NarX-NarL system in the regulation of biofilm components as well as *B. pseudomallei* host-associated survival. This study characterizes a conserved nitrate sensing system that is important in environmental and host-associated contexts and aims to bridge a gap between these two important *B. pseudomallei* lifestyles.

## Introduction

The ability of many bacteria, including bacilli and pseudomonads, to substitute nitrate as a terminal electron acceptor in oxygen-limited environments offers various benefits for niche adaptation at comparable free energy changes (1). Metabolism of N-oxides, inorganic ions comprised of oxygen and nitrogen, derived from exogenous sources in the environment or endogenous metabolic byproducts, preferentially follows the denitrification pathway (2). Exogenous N-oxides, such as nitrate (NO_3_^-^) and nitrite (NO_2_^-^), can be derived from anthropogenic environmental contamination or host innate immune cells during host-pathogen interactions. *Burkholderia pseudomallei*, an environmental saprophyte (3) and sapronotic disease agent (4), is an opportunistic pathogen that transitions between environments where nitrate metabolism can influence bacterial physiology and biofilm dynamics (5). Biofilms are comprised of extracellular polymeric substances that represent a protective matrix in which bacteria can reside; the formation and degradation of which is dependent on extracellular cues and intrinsic bacterial signal transduction mechanisms. *B. pseudomallei* 1026b, a clinical isolate (6), encodes a complete denitrification pathway and responds to exogenous nitrate to some extent by inhibiting biofilm formation and reducing intracellular cyclic di-GMP (5). The denitrification pathway in *B. pseudomallei* 1026b aligns with the conservation of denitrification genes among the β-proteobacteria, including *B. mallei* and *B. pseudomallei* (7). Utilizing a transposon library in *B. pseudomallei* 1026b, we have recently characterized a predicted two-component nitrate-sensing system (*narX*-*narL*) that negatively regulates biofilm formation and c-di-GMP production in the presence of nitrate (5). However, the genetic drivers of biofilm dynamics that link the signal transduction and metabolism of exogenous nitrate and c-di-GMP turnover in *B. pseudomallei* are poorly understood.

The NarX and NarL proteins comprise a conserved two-component regulatory system whereby NarX responds to nitrate and nitrite ligands to initiate autophosphorylation and activation of the NarL response-regulator receiver domain (8, 9). The *narX-narL* operon, as initially described in *Escherichia coli*, regulates transcription of gene clusters involved in fermentation and anaerobic respiration (10-13), and specifically activates *narGHJI* (membrane-associated nitrate reductase) and *narK* (nitrate/nitrite transporter) promoters to which it is adjacent (9) (Fig 1A). *Pseudomonas* and *Burkholderia* spp. contain a *narX-narL* system for nitrate and nitrite sensing (9), which directs hierarchical control of anaerobic respiration and general cellular physiology (14). The *narX-narL* system has been shown to regulate anaerobic metabolism in *P. stutzeri* (15) and *P. aeruginosa* (16), in which nitrate chemotaxis has recently been demonstrated and hypothesized to play a role in virulence gene regulation (17). The *narX and narL* genes have recently been implicated in the global regulation of *B. thailandensis* biosynthetic gene clusters via hierarchical control of host survival and anaerobic respiration genes (18). Saprophytic bacteria, including the *B. pseudomallei* complex, encode biosynthetic NRPS/PKS clusters that are important for survival in the rhizosphere but also may facilitate virulence in eukaryotic hosts (19). Although we have described nitrate sensing and metabolism in *B. pseudomallei* (5), we know little regarding the genetic regulation of biofilm physiology and the transcriptional profiles associated with nitrate respiration and host survival.

**Fig 1.**
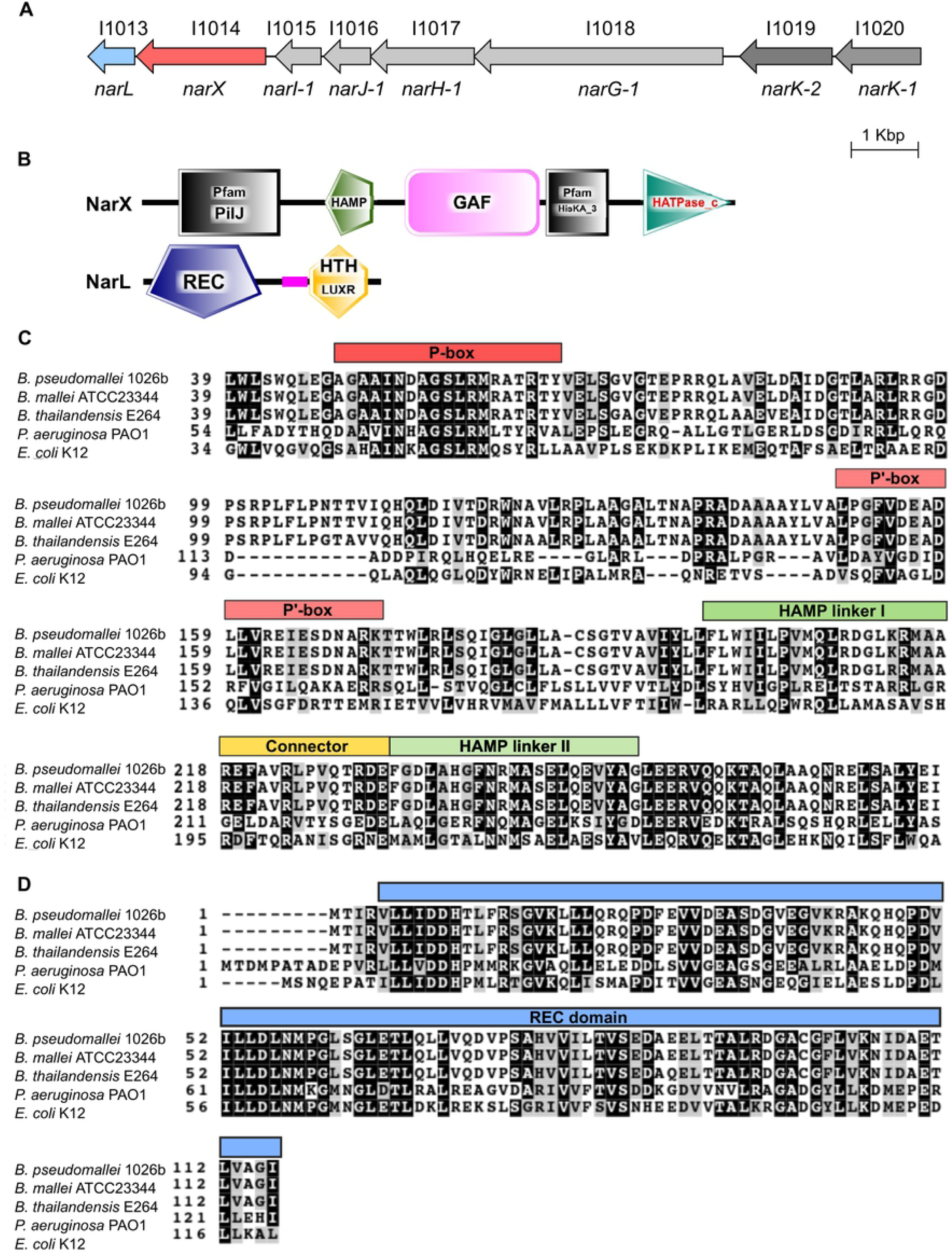
Orientation and conservation of the *narXL-narGHJI*_*1*_*-narK*_*2*_*narK*_*1*_ cluster. (A) Genomic orientation of the *narX-narL* regulatory system (red and blue, respectively), *narGHJI-1* dissimilatory nitrate reductase (light grey), and *narK-2* and *narK-1* nitrate/nitrite transporters (dark grey) in *B. pseudomallei* 1026b (coding sequences to scale). (B) Simple Modular Architecture Research Tool (SMART) protein domain analysis of NarX (Bp1026b_I1014) and NarL (Bp1026b_I1013). (C) Amino acid conservation of key residues in the sensory module of NarX (Bp1026b_I1014), including periplasmic domain sequences (P and P’ boxes) and HAMP (linker I, linker II, and connector) linker elements of *B. pseudomallei* 1026b, *B. mallei* ATCC 23344, *B. thailandensis* E264, *P. aeruginosa* PAO1, and *E. coli* K12. (D) Amino acid conservation of key residues of the receiver domain of NarL (Bp1026b_I1013) including the same organisms as above. Multiple sequence alignments were generated using Clustal Omega and visualized using BoxShade v3.2 where black boxes indicate identical residues and grey boxes indicate similar sequences.

*B. pseudomallei* is a facultative anaerobe that can adapt to oxygen tension in plant-associated rhizospheres (20) or to an intracellular lifestyle in animal immune cells (21); environments that are rich in N-oxides and reactive nitrogen intermediates (RNI). Given the broad environmental distribution of *B. pseudomallei* (22) and its ability to cause severe acute infections in people with no significant medical history (23, 24) or persistent chronic infections in immunocompromised individuals (25, 26), this organism possesses sensing systems to adapt to various niches and extracellular stressors (27). Analysis of the gene expression profile of *B. pseudomallei* during intracellular infection has revealed a temporal transcriptional adaptation that rapidly modulates bacterial metabolism and physiology inside macrophages (28). Although stimulated macrophages can inhibit intracellular growth of *B. pseudomallei* through RNI-dependent bactericidal mechanisms, *B. pseudomallei* has the ability to survive and replicate in phagocytes (29). One approach for identifying resistance mechanisms for RNI is to induce the expression of these mechanisms via sub-lethal levels of the stressor (30). We used concentrations of nitrate and nitrite that have been previously validated (31, 32), and which we have previously shown to be inhibitory to biofilm formation but not inhibitory to growth (5), to identify gene loci that are potentially involved in resistance to RNI and likely contribute to intracellular survival of *B. pseudomallei* in host cells.

Here, we describe global transcriptome profiling of gene expression in *B. pseudomallei* under nitrosative stress that is coordinated by the two-component nitrate-sensing system comprised of *narX* and *narL*. RNA-seq analysis of the Δ *narX* and Δ *narL* mutants revealed a global network of genes involved in nitrate/nitrite signal transduction and identified key elements that contribute to biofilm formation, virulence factors, antibiotic resistance markers, as well as the differential regulation of key natural product biosynthetic gene clusters. The nitrosative stress response is alleviated in the absence of either *narX* or *narL*. Additionally, we characterized the intracellular replication kinetics of *B. pseudomallei* lacking nitrate-sensing capabilities thereby linking *narX* and *narL* to pathogenicity and survival of this pathogen in the host. These results provide a framework for understanding biofilm inhibition mediated by the nitrosative stress response, and link secondary metabolite biosynthesis, stringent response, and intracellular survival to nitrate metabolism in *B. pseudomallei*.

## Methods

### Bacterial strains and growth conditions

*B. pseudomallei* 1026b, a clinical isolate from a human septicemic infection (33), was grown in Lysogeny Broth (LB) at 37°C with aeration as described previously (5), unless otherwise indicated. *E. coli* DH5α and RHO3 strains were grown in LB at 37°C with aeration. *B. pseudomallei* experiments were carried out in biosafety level 3 (BSL-3) in the Regional Biocontainment Laboratory at Colorado State University (5). For anaerobic growth, LB was supplemented with 0.75% glucose (LBG) and experiments were performed inside a 2.5 L anaerobic jar (AnaeroPack) using one sachet of Anaerobic Gas Generator (AnaeroPack). For experiments involving media with nitrate or nitrite, LB was supplemented with a final concentration of 10 mM sodium nitrate (Sigma) or 10 mM sodium nitrite (Matheson Coleman and Bell) as described previously (5), unless otherwise indicated.

### Mutant strain construction and complementation

In-frame deletion mutations were constructed as previously described (34). Briefly, *B. pseudomallei* Bp82 (35) (strain excluded from Select Agent regulations) genomic DNA was used as a template for PCR fragment generation in all experiments. DNA restriction enzymes were purchased from New England Biolabs. Primers and plasmids used for mutant strain construction and complementation are listed in S1 Table. In-frame deletion constructs were generated using Splicing Overlap Extension (SOE) PCR (36). Flanking sequences on both sides (∼1Kb each) of Bp1026b_I1014 (*narX*) and Bp1026b_I1013 (*narL*) were amplified from Bp82 template DNA, and spliced via amplification of a fragment excluding the gene coding regions. Spliced overlap fragments were cloned into pEXKm5 (37) and electroporated into *E. coli* RHO3 (37). *E. coli* RHO3 pEXKm5::Δ*narX* and pEXKm5::Δ*narL* were grown in LB supplemented with diaminopimelic acid (400 µg/mL) and kanamycin (35 µg/mL) at 37°C. Tri-parental mating with *B. pseudomallei* 1026b was facilitated with *E. coli* RHO3 pTNS3 grown in LB supplemented with ampicillin (100 µg/mL) and diaminopimelic acid (400 µg/mL). Transconjugants were selected for using kanamycin (1000 µg/mL) and X-Gluc (100 µg/mL) during screening. Merodiploids were resolved using yeast-tryptone (YT) media supplemented with 15% sucrose. In-frame deletion constructs were verified via internal and flanking primers (S1 Table). For complementation, full-length copies of Bp1026b_I1014 and Bp1026b_I1013 were cloned into pUC18T-mini-Tn*7*T-km-LAC for IPTG-inducible expression (38).

### Static biofilm and motility assays

Static biofilm and motility assays were performed as described previously (5, 39). All experiments were performed in LB supplemented with 10 mM sodium nitrate or 10 mM sodium nitrite unless otherwise specified, grown at 37°C, in either aerobic or anaerobic conditions. Biofilms were assayed in replicates of six individual wells of 96-well polystyrene plates (Nunc™ Microwell™ 96-well microplates #243656, Thermo Scientific) and processed as previously described (5). Motility was measured by inoculating strains of interest into 0.3% semisolid LB agar, supplemented with sodium nitrate as indicated, and measuring visible diameter of bacterial spread over the specified time.

### Nitrite ion measurement

Nitrite ion (NO_2_^-^) from bacterial cultures was measured using the Griess Reagent system (Promega) following the protocol recommended by the manufacturer. Briefly, a nitrite standard reference curve was generated and included on each 96-well plate (Nunc) used for experimental samples. The Griess reaction was performed using room-temperature sulfanilamide solution and NED solution and the resulting azo compound density was measured at OD_550_. These experiments were performed in aerobic as well as anaerobic conditions with wild type, Δ *narX*, and Δ *narL* mutant strains grown statically for 24 hours at 37°C in biological triplicates and technical triplicates. Aerobic cultures were cultivated in LB media supplemented with 10 mM NaNO_3_ and anaerobic cultures were cultivated in LBG media (LB 0.75% glucose) and supplemented with 25 mM NaNO_3_.

### RNA isolation and RNA-seq library preparation

Total RNA was isolated from static-growth cultures as described previously (5), with a few modifications. LB media was used due to previous observations of pellicle biofilm formation, in either aerobic or anaerobic conditions as indicated. Pellicle biofilms were grown in six-well Costar polysterene plates (Corning) for 24 hours at 37°C, at which point 1.5 mL of culture samples were collected and resuspended in RNAprotect Bacteria Reagent (Qiagen) and then QIAzol Lysis Reagent (Qiagen) before storage at −80°C. RNA samples were purified and depleted of genomic DNA as described previously (5), before depletion of ribosomal RNA with Ribo-Zero rRNA Removal Kit for bacteria (Illumina) and purification using magnetic beads (AMPure). RNA-seq libraries of cDNA were generated using ScriptSeq™ Complete v2 RNA-seq Library Preparation Kit (Illumina) and purified using Monarch DNA cleanup kit (New England Biolabs). Unique barcodes were added to each sample library using ScriptSeq™ Index PCR Primers (Illumina). Libraries were analyzed on a Tapestation using HS D1000 tapes and reagents (Agilent) to determine average sizes and concentrations of the libraries. Size and molarity estimates were used to pool all libraries in equimolar concentrations. Final quality control and library quantification analyses were completed at the Microbiology, Immunology, and Pathology Next Generation Sequencing Core Facility (MIP NGS Illumina Core) at Colorado State University.

### Illumina sequencing and differential expression quantification

A NextSeq run was completed on the pooled libraries using the NextSeq 500 hi-output v2 75-cycle kit and Buffer Cartridge (Illumina). Sequence files were downloaded from the MIP NGS server, de-multiplexed according to index primers, and converted to fastq files before initial quality control using FastQC (40). Adapter sequences were trimmed using Trimmomatic (ILLUMINACLIP:TruSeq3-SE:2:30:10 LEADING:3 TRAILING:3 SLIDINGWINDOW:4:15 MINLEN:72) before another quality control round using FastQC. Bowtie2 was used to align sequencing reads to the reference genome GCF_000260515.1_ASM26051v1 (NCBI) and TopHat was used for transcriptome assembly. HTseq-count (version 0.11.0) was used to count accepted hits before the DEseq2 (version 1.20.0) (41) package was employed in R (version 3.6.1) for comprehensive differential expression analysis. Raw read count coverage values were used to compare the differential gene expression between temperature treatments, mutants, and untreated controls. Using a negative binomial distribution to estimate variance and a Bayesian approach for variance shrinkage, the DEseq2 package produced logarithmic fold-change values between the conditions tested. Wald tests were used to calculate p-value and the Benjamini-Hochberg multiple testing correction was used to correct for the false discovery rate.

### Gene expression and quantitative real-time PCR

Genomic DNA-depleted RNA samples, isolated in quadruplicate, were pooled and cDNA was synthesized using 1 µg total RNA, using the Transcriptor First Strand cDNA Synthesis kit (Roche). Primers (S1 Table) for Bp1026b_I1018 (*narG-1*), Bp1026b_II1965 (CPS III) Bp1026b_I1913 (*relA*), Bp1026b_II1245 (bactobolin), Bp1026b_II1353 (syrbactin), and Bp1026b_I0189 (*hcp-1*), were designed using the PrimerQuest tool (IDT DNA Technologies), and primer efficiencies were calculated using dilutions of cDNA samples. The 23S rRNA reference gene (42) was used as a housekeeping control for normalization (43), due to its consistent expression profile for all cDNA samples even after attempted ribodepletion. qRT-PCR was performed using 10 ng total cDNA and the same cycling conditions described previously (5). Relative transcript abundance was measured using the Pfaffl method (44).

### Infection of murine macrophage cells

Murine macrophage cells (RAW 264.7 cell line, ATCC) were propagated in Dulbecco’s Modified Eagle Medium (DMEM, Gibco) supplemented with 10% fetal bovine serum and growth at 37°C with 5% CO_2_ and 80-90% relative humidity in T-75 flasks (Corning). *B. pseudomallei* 1026b strains were grown overnight at 37°C with aeration to stationary phase and diluted to an appropriate OD_600_ for an infection inoculum of ∼1×10^6^ CFU/mL. RAW cells were seeded in 12-well cell culture dishes (Corning) at a density of ∼5×10^5^ cells. Bacteria were resuspended in DMEM and added to RAW cells at a MOI of 5 and incubated at 37°C for 1 hour. RAW cells were washed with 1X PBS and culture medium was replaced with fresh medium supplemented with kanamycin (750 µg/mL) before a 2-hour incubation at 37°C to kill extracellular bacteria. RAW cells were washed and lysed in 1X PBS containing 0.2% Triton X-100 for 2.5 min and serially diluted and plated on LB agar to quantify intracellular bacteria. Survival of the wild type, Δ*narX*, and Δ*narL* strains were analyzed in macrophage using the same infection procedure above. RAW 264.7 cells were incubated with bacteria at MOI of 5 for 20 hours at 37°C and sampled at 6h and 20h. Cells were washed, lysed and serially diluted to plate for CFU/mL as described above.

## Results

### NarX-NarL comprise a nitrate-sensing two-component system in *B. pseudomallei*

The NarX-NarL two-component system (TCS) has been described extensively in γ-proteobacteria (9), in which it was discovered to regulate nitrate respiration in coordination with an additional NarQ-NarP TCS depending on environmental nitrate availability (45-47). In β-proteobacteria, *Burkholderia* and *Ralstonia* spp. encode only the NarX-NarL system, a feature shared with Pseudomonadaceae in the γ-proteobacteria clade but not Neisseriaceae (also β-proteobacteria) (9). The NarX-NarL TCS is often part of a larger regulon including the dissimilatory nitrate reductase NarGHJI and the transporters/permeases NarK-1 and NarK-2, as is the case in *B. pseudomallei* 1026b (Fig 1A). Analysis of this system via the SMART algorithm revealed a predicted HAMP domain in NarX and a predicted REC domain in NarL (Fig 1B), indictative of two-component signal transduction capabilities. NarX (*E. coli*) is a sensory histidine kinase with a periplasmic sensor domain that is dimeric when bound to its ligand, nitrate (NO_3_^-^) (48). The NarX (*E. coli*) sensory module (Fig 1C) contains conserved periplasmic domains (P-box and P’-box) that are necessary for ligand binding and response, yet only Gly^51^ and Met^55^ (numbering with respect to NarX_Ec_K12_) have been found to be invariant (9). The HAMP linker-connector-linker domains are common to sensor kinases and are required for signal transduction (49, 50). The *B. pseudomallei* complex (Bpc) which includes *B. mallei* ATCC 23344, an obligate intracellular pathogen, and *B. thailandensis* E264, a non-pathogenic soil saprophyte, share conserved NarX sensory module (Fig 1C) and NarL receiver domain (Fig 1D) sequences with *P. aeruginosa* PAO1 and *E. coli* K12. Additionally, the complete *narX-narL-narGHJI*_*1*_*-narK*_*1*_*-narK*_*2*_ regulon is conserved in its entirety in *B. mallei* ATCC 23344 based on *in silico* sequence alignment (S1 Fig). Collectively, these bioinformatics analyses suggest that the NarX-NarL system functions as an archetypal signal-response regulatory system in *B. pseudomallei* 1026b, and that organisms across the Bpc share this functional genomic element.

### NarX and NarL respond to nitrate but not nitrite to inhibit biofilm formation

Previously, we have shown that both sodium nitrate and sodium nitrite inhibit *B. pseudomallei* 1026b biofilm formation in a dose-dependent fashion (5). Five genes, *narX, narL, narH*_*1*_, *narG*_*1*_, and *narK*_*2*_, were implicated in the regulation of biofilm dynamics in response to exogenous nitrate and nitrite (5). Deletion mutants Δ*narX* and Δ*narL* and the wild type were grown statically in LB supplemented with either 10 mM NaNO_3_ or 10 mM NaNO_2_ for 24h (Fig 2A). Biofilm inhibition via nitrate and nitrite supplementation was observed for the wild type while both Δ*narX* and Δ*narL* mutants were resistant to nitrate biofilm inhibition but not nitrite. Nitrite inhibited biofilm formation in the nitrate sensing-deficient mutants at similar levels to wild type (Fig 2A). Nitrate-mediated biofilm inhibition of Δ*narX* and Δ*narL* mutants was restored by complementation of Bp1026b_I1014 (*narX*) and Bp1026b_I1013 (*narL*)) with IPTG-induction and was comparable to the wild type (Fig 2B). Consistent with our previous transposon insertional mutants (5), both components of the NarX-NarL system respond to nitrate by biofilm inhibition, however this system is not similarly affected by nitrite at the concentration tested. These results suggest that the NarX-NarL system has specificity for nitrate sensing in the regulation of biofilm dynamics in *B. pseudomallei* 1026b.

**Fig 2.**
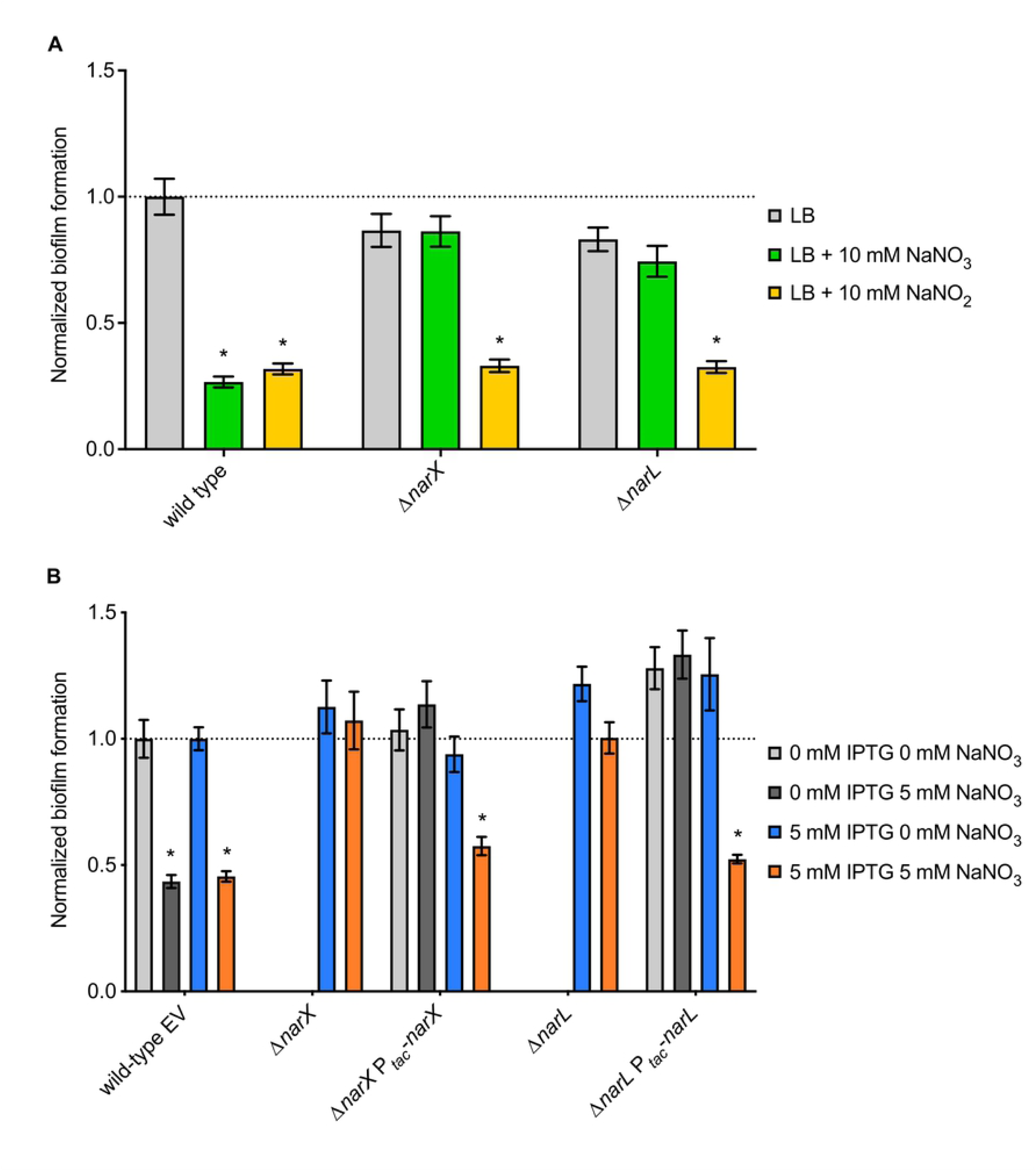
Biofilm formation of *B. pseudomallei* Δ*narX, and* Δ*narL* strains and their complements as compared to the wild type. (A) Wild type, Δ*narX*, and Δ*narL* biofilms were grown in LB media (grey) supplemented with either 10 mM NaNO_3_ (green) or 10 mM NaNO_2_ (yellow) at 37°C for 24 h. (B) Complements of Δ*narX* and Δ*narL* mutants were grown in LB media (light grey) supplemented with 5 mM NaNO_3_ (dark grey), 5 mM IPTG (blue), or 5 mM IPTG and 5 mM NaNO_3_ (orange), and compared to wild-type *B. pseudomallei* 1026b containing an empty complementation vector (EV). Asterisks indicate significant differences (* = p < 0.0001) calculated with an unpaired Student’s T-test using the Bonferonni method to account for multiple comparisons (n = 12).

### Nitrite, but not nitrate, suppresses anaerobic biofilm growth in *B. pseudomallei*

To examine the effects of nitrate and nitrite on oxygen-deprived bacterial cells, as commonly found in tissue-associated biofilm (51) or intracellular infections (52), we adapted an anaerobic biofilm model for *B. pseudomallei* (53). Static biofilm growth was initiated in an oxygen-deprived anaerobic jar with the addition of an added carbon source (0.75% glucose) and enhanced via supplemented nitrate, which increased anaerobic biofilm formation starting at 10 mM (Fig 3A). In contrast to the beneficial effect of nitrate, the addition of sodium nitrite had an inhibitory effect on anaerobic biofilm growth (Fig 3B). Significant biofilm growth defects were observed starting with the addition of 10 mM NaNO_2_, while subsequent concentration increasingly favored growth in NaNO_3_-supplemented media (Fig 3B). The observed cessation of growth in the nitrite treatment corresponds to a similar phenotype involving mycobacterial growth repression due to endogenous nitrite accumulation (52). Interestingly, the addition of exogenous nitrate for anaerobic swim motility assays showed a robust increase in flagellar motility in response to nitrate sensing, considering that absence of both *narX* and *narL* inhibited swimming even with nitrate present (Fig 3C).These results suggest that *B. pseudomallei* uses nitrate but not nitrite as an alternative terminal electron acceptor during anaerobic biofilm growth, and that nitrite may inhibit anaerobic growth, suggesting hierarchical control during the shift to anaerobiosis.

**Fig 3.**
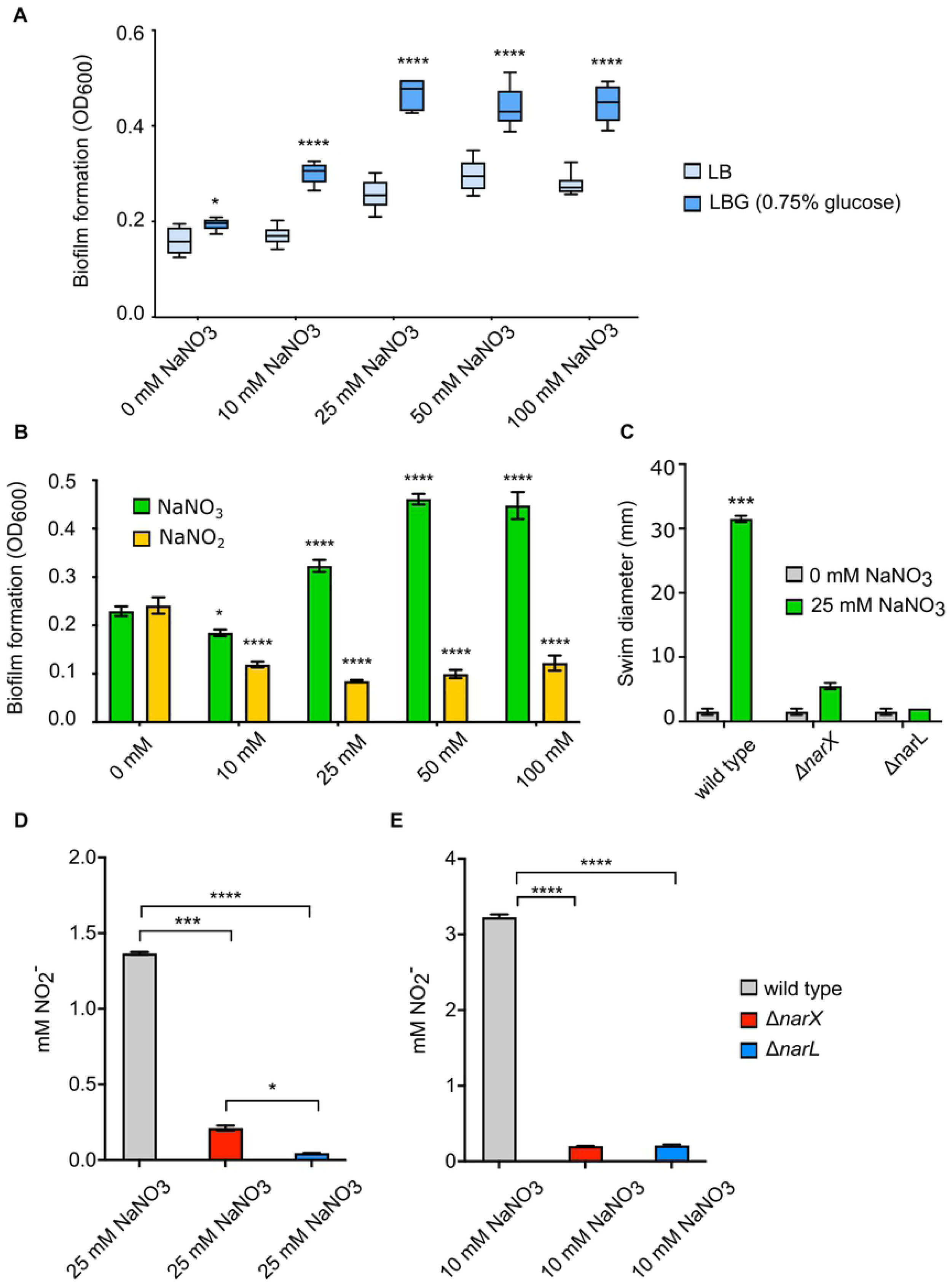
Anaerobic biofilm growth of *B. pseudomallei* utilizing an alternative terminal electron acceptor. (A) Static growth of wild-type *B. pseudomallei* was measured after 24 hours of anaerobic growth at 37°C with increasing concentrations (0, 10, 25, 50, and 100 mM) of NaNO_3_ in LB media (light blue) or LB media supplemented with 0.75% glucose (dark blue). Asterisks indicate significant differences (* = p < 0.05, **** = p <0.0001) calculated with an unpaired Student’s T-test using the Bonferonni method to account for multiple comparisons (n = 12). (B) Static growth of wild-type *B. pseudomallei* was measured after 24 hours of anaerobic growth at 37°C grown in LBG (LB 0.75% glucose) media supplemented with either NaNO_3_ (green) or NaNO_2_ (yellow) at increasing concentrations (0, 10, 25, 50, and 100 mM). Asterisks indicate significant differences (* = p < 0.05, **** = p <0.0001) calculated using a Dunnett’s multiple comparison 2-way ANOVA test. (C) Swimming motility of wild type, Δ*narX*, and Δ*narL* strains in 0.3% semi-solid LBG agar (grey) supplemented with 25 mM NaNO_3_ (green) and incubated at 37°C for 24 hours. Asterisks indicate significant differences (*** = p <0.001) calculated with an unpaired Student’s T-test using the Bonferonni method to account for multiple comparisons (n = 6). Griess reaction results measuring nitrite ion produced by wild type (gray), Δ*narX* (red), and Δ*narL* (blue) strains anaerobically (D) after growth in LBG with 25 mM NaNO_3_ or aerobically (E) after growth in LB with 10 mM NaNO_3_ at 37°C for 24 hours. Asterisks indicate significant differences (* = p < 0.05, *** = p < 0.001, **** = p <0.0001) calculated with an unpaired Student’s T-test using the Bonferonni method to account for multiple comparisons (n = 12).

To analyze relative function of the primary nitrate reductase in *B. pseudomallei* under both aerobic and anaerobic conditions, we next examined the production of nitrite in culture media using the Griess test. Using Griess reagent, which enables colorimetric quantification of nitrite ion in solution, we measured 1.4 mM NO_2_ and 3 mM NO_2_ in anaerobic (Fig 3D) and aerobic (Fig 3E) culture conditions, respectively. For Griess assays, we supplemented LB medium with 10 mM NaNO_3_ and 25 mM NaNO_3_ in aerobic and anaerobic conditions, respectively, to account for both biofilm inhibition and anaerobic growth models. In both conditions, absence of either *narX* or *narL* significantly inhibited the production of nitrite ion in solution (Figs 3D and 3E), indicating that the predicted nitrate-sensing two-component system activates nitrate-nitrite respiration via the primary nitrate reductase in both aerobic and anaerobic conditions. Altogether, these results demonstrate a disparity between exogenous nitrate or nitrite supplementation regarding biofilm growth in *B. pseudomallei*; while both nitrate and nitrite inhibit aerobic biofilm growth, nitrite suppresses anaerobic biofilm growth. Both components of the nitrate-sensing system, *narX* and *narL*, respond to exogenous nitrate and nitrite in a similar fashion by facilitating nitrate-nitrite respiration, motility, and biofilm inhibition.

### Analyses of global responses to nitrate/nitrite-mediated biofilm inhibition reveals divergent responses

Because of the observed disparities between nitrite-mediated biofilm inhibition in Δ*narX* and Δ*narL* strains (Fig 2A), as well as our previous observations involving decreased cyclic di-GMP production in response to nitrate (5), we next aimed to characterize the global transcriptional response to nitrate/nitrite sensing in *B. pseudomallei* 1026b. To assess the impacts of nitrosative stress on the *B. pseudomallei* transcriptome, wild type, Δ*narX*, and Δ*narL* strains were grown statically in LB media or supplemented with either 10 mM NaNO_3_ or 10 mM NaNO_2_, in biological quadruplicates. RNA from technical triplicates were collected and pooled on separate days for each biological sample to minimize batch effect, then treated with DNase I, depleted of ribosomal RNA, chemically fragmented, and reverse transcribed into indexed cDNA libraries (Fig S2). Illumina sequencing using a NextSeq 500 hi-output flow cell, followed by read quality assessment and subsequent mapping and alignment to the *B. pseudomallei* 1026b genome, and statistical analysis of differentially regulated transcripts via DESeq2 revealed divergent transcriptome datasets (Fig 4). The minimal genetic relatedness between these populations of cells responding to differing environmental conditions is supported by the hierarchical clustering and ordination of rlog-transformed data for each transcript Z-score that is linked by sample and treatment condition (Figs 4A and 4B). When comparing wild-type cultures grown in LB supplemented with or without 10 mM NaNO_3_, principal component analysis reveals unrelated populations with 77% of variance explained by one principal component (Fig 4C). Similar results were observed when comparing wild-type cultures grown in LB supplemented with or without 10 mM NaNO_2_, showing that the first principal component explains 74% of the variance (Fig 4D). However, when comparing the transcriptome data to account for nitrate and nitrite effects on wild type, most differentially regulated transcripts are affected similarly by nitrate and nitrite (Fig S3). Venn diagrams display 181 up-regulated genes (Fig S3A) with fold changes greater than two and 203 down-regulated genes (Fig S3D) with fold changes less than two and false discovery thresholds below *q* < 0.01 that share expression profiles between nitrate- and nitrite-treated sample groups. These analyses on the global transcriptome datasets suggest that although substantial differences exist in the response to growth supplemented with either nitrate or nitrite, these genetic differences are mitigated when comparing nitrate and nitrite treatments to each other.

**Fig 4.**
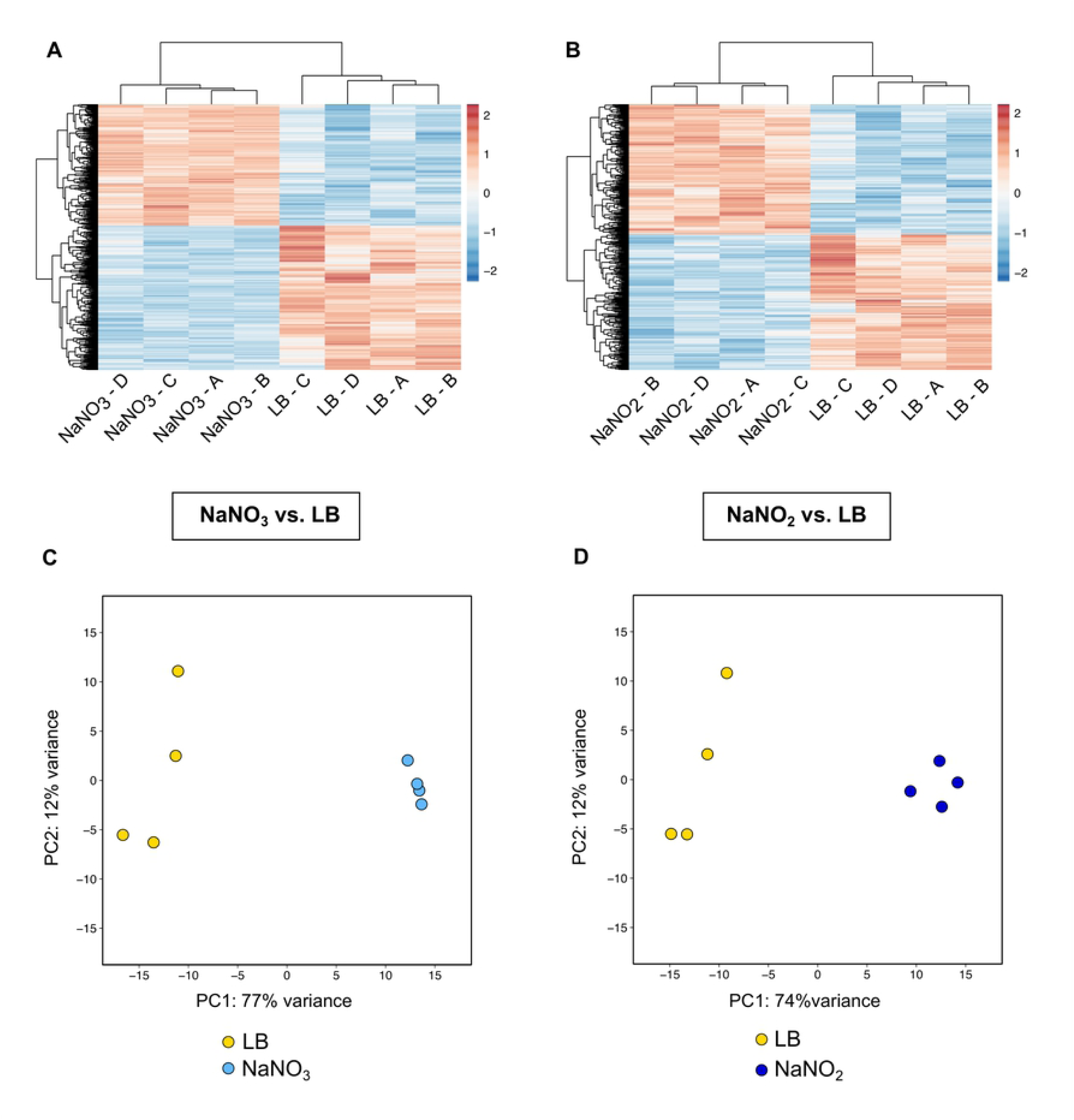
Clustering analyses reveal divergent datasets in the differential expression patterns among wild type samples in both NO_3_^-^ and NO_2_^-^ treatment conditions. Z-score analysis of all differentially regulated transcripts for NaNO_3_ vs. LB (A) and for NaNO_2_ vs. LB (B). Principal component analysis of entire datasets for NaNO_3_ vs. LB (C) and for NaNO_2_ vs. LB (D). Hierarchical clustering of all significantly expressed transcripts are presented in panels A and B and principal component analyses represent the complete transcriptomes of all samples as depicted in panels C and D. Principal components were calculated using the regularized log transformation of raw transcript count data in the DESeq2 package.

### Differential expression profiles reveal key metabolic processes, biofilm components, respiratory pathways, and biosynthetic clusters that are globally regulated in response to nitrate and nitrite

To analyze the global gene expression profiles of wild-type cells grown in the presence of nitrate or nitrite, Illumina single-end sequencing reads using the DESeq2 modeling and statistical package for RNA-seq data were analyzed (41). Differential expression analysis was performed using pair-wise comparisons across the three strains (wild type, Δ*narX*, and Δ*narL*) and the three treatment conditions (LB, LB 10 mM NaNO_3_, and LB 10 mM NaNO_2_), with four biological samples per group. In total, 36 samples were assembled into cDNA libraries, indexed with individual barcodes, and pooled for NextSeq Illumina Sequencing, resulting in 408.8 M reads passing initial filtering. Following further filtering, trimming, mapping, and alignments, transcript count matrices for each sample were compiled using the HTseq Python package (54). Un-normalized count matrices were then imported into RStudio and the DESeq2 differential expression pipeline with thresholds and outputs specified in R. For significantly differentially expressed genes, we set thresholds of log_2_ Fold Change < −1 or > 1 and adjusted *p*-value < 0.001. Overall, 274 up-regulated genes and 316 down-regulated genes were observed in the nitrate-supplemented condition versus LB for wild type (Fig 5A). 237 genes were up-regulated and 271 were down-regulated in the nitrite condition versus LB for the wild type as well (Fig 5B). However, when comparing the Δ*narX* mutant to the wild type in the nitrate supplemented condition, the trends are reversed (Fig 5C), further supporting that nitrate sensing is required for biofilm inhibitory phenotype. A similar trend was observed in the Δ*narL* mutant under the same conditions (data not shown), which is expected considering the overlap of gene regulation of both mutants (Figs S3B and S3E). We identified key differences and similarities regarding the effects of nitrate and nitrite on the differential regulation of metabolic, respiratory, biofilm, virulence, and biosynthetic gene clusters in *B. pseudomallei* 1026b as detailed below (Fig 6). Given the significant overlap in transcripts regulated similarly by both nitrate and nitrite, we explored the genome-wide regulation of gene clusters of proximal genes with the same differential regulation and visualized the effect of exogenous nitrate on the *B. pseudomallei* 1026b genome as detailed below (Fig 6). Using the Webserver for Position Related data analysis of gene Expression in Prokaryotes (WoPPER) (55), we considered the transcriptional regulation of exogenous nitrate to identify differentially expressed regions on each chromosome of *B. pseudomallei* 1026b. By identifying and mapping physically contiguous clusters of genes along both chromosomes that respond similarly to nitrate, we can provide a visual validation of the differential expression statistical output provided by DESeq2. While there is no visual difference between the regulation profiles among both chromosomes, this graphical representation of the gene expression data set highlights important biofilm-associated clusters, antibiotic resistance-associated clusters, and interestingly, the confinement of secondary metabolite biosynthetic gene clusters on chromosome II. In total, WoPPER analysis identified 27 clusters on chromosome I (S2 Table) and 21 clusters on chromosome II (S3 Table) as differentially regulated in response to exogenous nitrate. The following sections explore the trends described above (Figs 5 – 8) in more detail, considering distinct loci (S4 – S7 Tables) in relation to their metabolic and biosynthetic functions.

**Fig 5.**
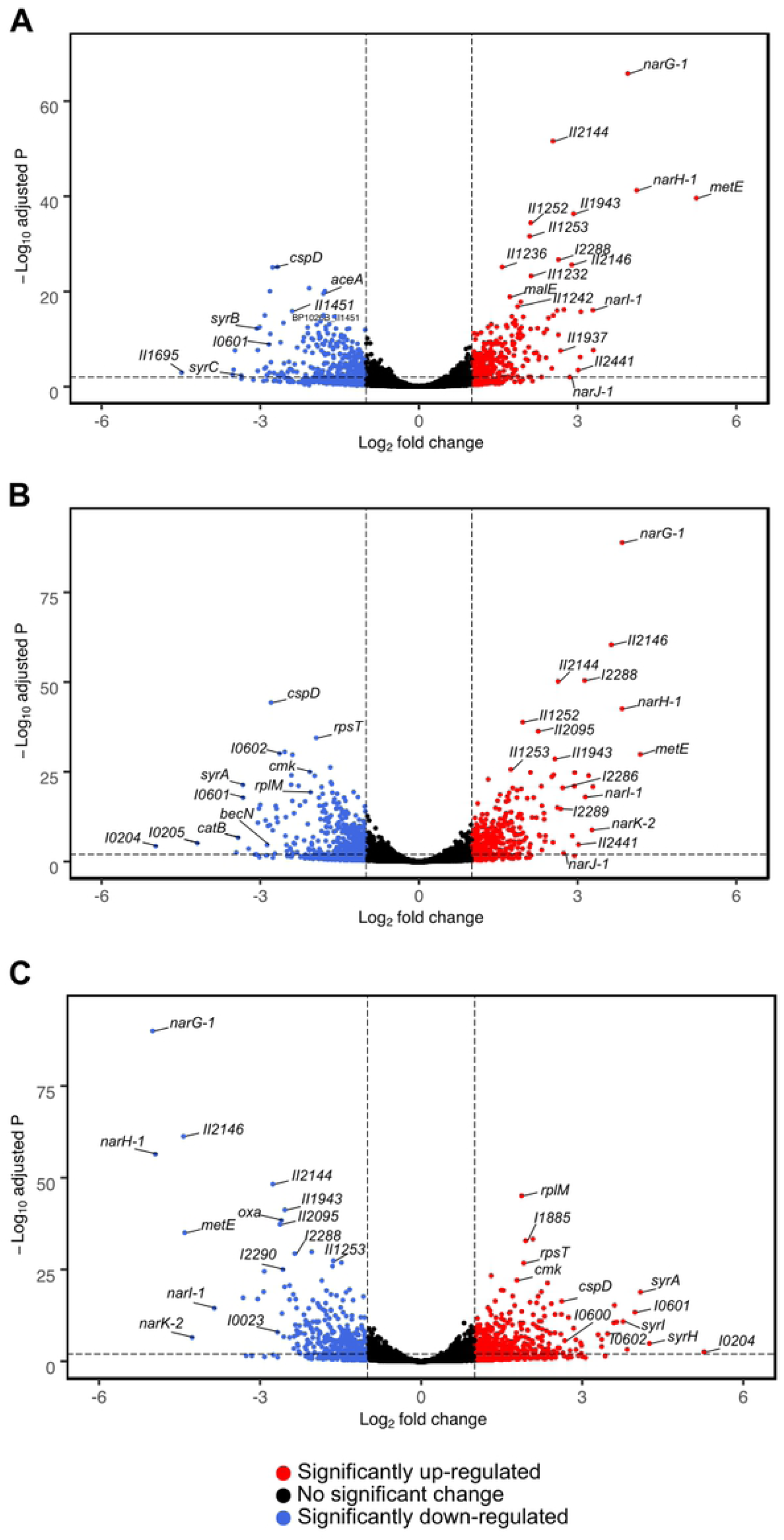
Trends in transcript fold changes in relation to statistical significance for comparably regulated datasets under nitrosative stress are reversed in the Δ*narX* mutant. NO3 ^-^ vs. LB (A) and NO_2_ ^-^ vs. LB (B), and Δ*narX* vs. wild type in the NO_3_ ^-^ condition (C). Transcripts are depicted as colored circles; unchanged or middle expression (black), upregulated or high expression (red), downregulated or low expression (blue). Log_2_ fold change values are distributed along the X-axes and Log_10_ adjusted p-values (as determined by the Wald test) are distributed along the Y-axes.

**Fig 6.**
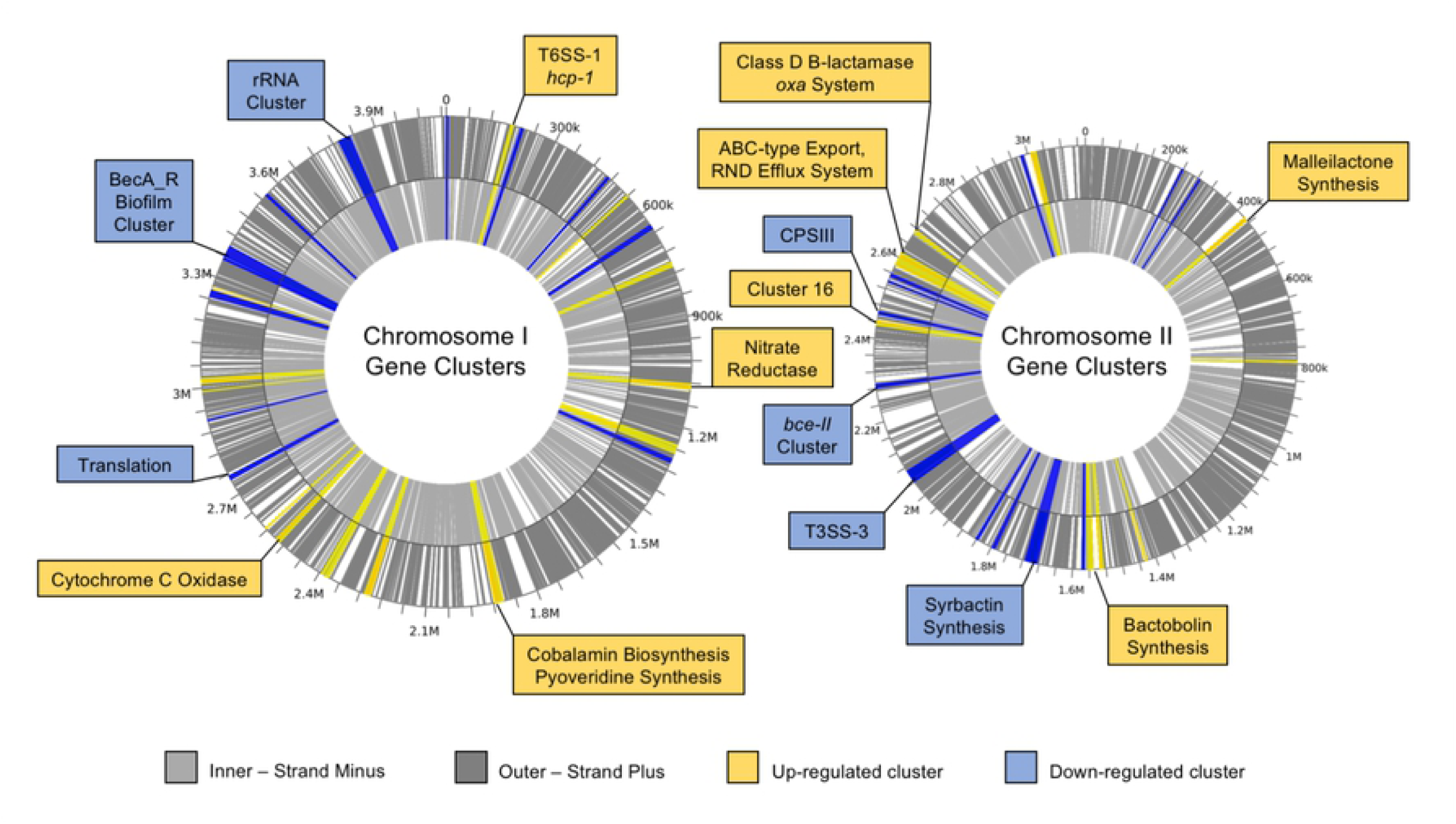
Differentially regulated transcripts comprise clusters of genes distributed across both chromosomes in *B. pseudomallei* 1026b. WoPPER analysis reveals biofilm-associated gene clusters, secondary metabolic biosynthetic clusters, general metabolism and respiration, and virulence-associated clusters are differentially regulated in response to nitrate treatment. Up-regulated clusters are highlighted in yellow, and down-regulated clusters in blue. Inner and outer circles represent positive (dark gray) or negative (light gray) strands of DNA. For complete transcriptional analysis of each cluster see Tables S2 and S3.

### Nitrate metabolism

Not surprisingly, among the most highly expressed transcripts in both nitrate and nitrite treatment groups compared to LB were genes in the *narXL-narGHJI-narK*_*2*_*-narK*_*1*_ regulon. Bp1026b_I1015 (*narI-1*), I1016 (*narJ-1*), I1017 (*narH-1*), I1018 (*narG-1*), I1019 (*narK-2*), and I1020 (*narK-1*) were collectively up-regulated at mean fold change of 9.7 in the nitrate treatment group, and at mean fold change of 10.5 in the nitrite group. Among these loci, the α- and β-subunits of the dissimilatory nitrate reductase *narG-1*, and *narH-1*, respectively, were the most highly and significantly expressed in the nitrate and nitrite groups (S4 Table, S6 Table). Noticeably, only loci in the *narXL-narGHJI-narK*_*2*_*-narK*_*1*_ regulon were significantly regulated in response to either nitrate or nitrate, omitting loci on chromosome II and the duplicate dissimilatory nitrate reductase *narGHJI-2*. Genes encoding assimilatory nitrate and nitrite reductases, nitrite reductases, nitric oxide, and nitrous oxide reductases were all absent from our expression data. The absence of other nitrate metabolism genes in this analysis suggests that the normoxic experimental conditions did not stimulate anaerobic respiration and potentially implicate the nar*X-narL* regulon in nitrate-nitrite respiration.

Bp1026b_I1014 (*narX*) was noticeably down-regulated in both treatment groups; −2.5-fold in the nitrate group and −2.9-fold in the nitrite group. This seeming discrepancy can be explained by the fact that the snapshot into global regulatory mechanisms provided by these RNA-seq data are reflective of a 24-hour time-point, at which point nitrate sensing has ceased and nitrate metabolism is well underway. The response regulator and transcription factor Bp1026b_I1013 (*narL*) was also absent from this differential expression analysis. Bp1026b_0861 (*mobA*), encoding for the final molybdoenzyme responsible for generating the molybdenum guanine dinucleotide cofactor required for nitrate reductase function (56) was also down-regulated in both nitrate and nitrite treatment groups; −2.9-fold and −2.2-fold, respectively. The down-regulation of key genes associated with activation of nitrate reduction can be explained by the sampling time after prolonged exposure to the nitrosative treatments. Altogether, under the conditions tested in these transcriptomic analyses, the primary nitrate reductase, which is activated at 10 mM NO_3_ ^-^ or above (57), along with the *narX-narL* system and the *narK* extrusion genes are the key elements of nitrate metabolism in *B. pseudomallei*.

### General metabolism

In addition to nitrate metabolism, several transcripts associated with amino acid and energy metabolism were identified as being significantly regulated in response to nitrate and nitrite via differential expression. The highest and most significantly expressed transcript for both treatment groups, Bp1026b_I0767 (*metE*), (Figs 5A, 5B) is a cobalamin-independent methionine synthase that produces methionine from homocysteine (58, 59). *metE* was up-regulated 18.3-fold in the nitrate group and 37.9-fold in the nitrite group. *metE* has been shown to be strongly induced in *R. solanacearum* during plant cell infection (54) and implicated in homocysteine turnover in *S. typhimurium* during infection where other methionine metabolism enzymes are candidate RNI resistance genes (30, 60). In addition to methionine metabolism, sulfate metabolic loci (Bp1026b_I1794 – I1798) were up-regulated a mean 3.1-fold and 4.0-fold in the nitrate and nitrite treatment groups, respectively. A genetic cluster containing 12 loci involved propionate metabolism (Bp1026b_II1934 – II1945) was up-regulated by a mean 4.5-fold and 5.0-fold in the nitrate and nitrite treatment groups, respectively. The gene cluster containing the predicted propionate metabolism loci is also annotated as a non-ribosomal peptide synthase cluster (Cluster 16) of unknown biosynthetic functionality in both *B. pseudomallei* and *B. thailandensis* (61, 62), with Bp1026b_II1945 in *B. pseudomallei* 1026b annotated as a predicted peptide synthase. However, two loci within this cluster are analogous to the *mhpE* aldolase (II1937) and *mhpF* dehydrogenase (II1938) loci in *E. coli* with putative aromatic cleavage function (63). Propionate is an important carbon source in proteobacteria that is intricately linked to central metabolism and the TCA cycle, as it can be converted to pyruvate via *prpB* and the PrpR sigma factor (64).

Correspondingly, Bp1026b_II0229 (*prpB*) and Bp1026b_II0228 (*prpR*) were both down-regulated 2- to 3-fold in in both the nitrate and nitrite treatment groups. Bp1026b_I1208 (*aceA*, isocitrate lyase) and Bp1026b_I1204 (*aceB*, malate synthase), genes involved in the glyoxylate cycle of central metabolism and carbohydrate synthesis, were also significantly down-regulated 2- to 4-fold in both treatment groups. Collectively, these data suggest dysregulation in the TCA cycle due to nitrosative stress and present intriguing possibilities regarding bacterial defense from RNI using essential metabolic enzymes such as *metE*. These data also complement a recent transcriptomic analysis comparing a *B. pseudomallei* Δ *gvmR* (globally acting virulence and metabolism regulator) mutant relative to the wild type (65). In our analysis, Bp1026b_I0113 (*gvmR*) is significantly down-regulated −2.7-fold in both the nitrate and nitrite treatment groups; however, a pairwise comparison with Δ*narX* versus the wild type under nitrate treatment reverses this trend and *gvmR* is up-regulated 2-fold. Thus, the general nitrosative stress response in *B. pseudomallei* involves significant shifts in central metabolic processes as well as differential regulation of several virulence determinants.

### Virulence-associated genes

Several antigens and secretion systems were differentially regulated in response to nitrate and nitrite treatment in *B. pseudomallei* 1026b. Notably, the type VI secretion system (T6SS) gene *hcp-1* from T6SS cluster 1 was significantly upregulated 2.3-fold in the nitrate condition and 2.5-fold in the nitrite condition. Organisms in the Bpc, including *B. pseudomallei, B. mallei*, and *B. thailandensis* variably encode six separate T6SS clusters, with *B. pseudomallei* having the coding capacity for all six (66). Of these six, T6SS-1 has been shown to be a virulence determinant in *B. mallei* (66), and hemolysin co-regulated protein-1 (*hcp-1*) has been shown to be important for eukaryotic cell replication and multinucleated giant cell (MNGC) formation in *B. pseudomallei* (67). However, T6SS eukaryotic effectors such as Hcp-1 are tightly regulated for infection and are usually not expressed or produced in laboratory growth conditions (67, 68). While *hcp-1* expression has been shown to be negatively regulated by zinc and iron (68), our results indicate positive regulation by nitrate and nitrite that requires an intact NarX-NarL sensing system (*hcp-1* is down-regulated in Δ*narX* under nitrosative stress). In contrast, expression of genes in the type III secretion system (T3SS) cluster 3 was significantly reduced in the nitrate and nitrite conditions. Comprised of 37 genes, T3SS-3 is a major virulence determinant in *B. pseudomallei* (69) and has been shown to participate in intracellular spread and actin-based motility in macrophage-like cells (70, 71). Our analysis identified three genes within this cluster, Bp1026b_II1628 (*bipB*), II1643 (*bsaM*), and II1644 (*bsaN*) that were significantly down-regulated at a mean fold change ratio of −4.5 in the nitrate condition (S5 Table) and −5.8 in the nitrite condition (S7 Table). *bsaN* has been shown to encode a positive transcriptional regulator of the T3SS-3 cluster in *B. pseudomallei* that is down-regulated in response to arabinose (72). In a similarly indirect fashion, growth with supplemental nitrate and nitrite suggest that exogenous N-oxides have implications for *B. pseudomallei*, in this inhibiting T3SS-3-mediated pathogenesis.

Again, these changes are dependent on a functional NarX-NarL system, as four T3SS-3 loci in Δ*narX* were up-regulated at a mean 4.2-fold compared to the wild type in the nitrate condition. In addition to these secretion system-associated gene loci, our analysis identified two loci within the capsular polysaccharide 1 (CPS I) cluster as being positively regulated in both nitrate and nitrite treatment groups. Bp1026b_I0505 (*wzt*), a putative ATP-binding ABC transporter capsular polysaccharide export protein, and Bp1026b_I0503 (*wcbD*), capsular polysaccharide export system inner membrane protein, were up-regulated at a mean fold change ratio of 2.0 in the nitrate supplemented condition and 2.3 in the nitrite supplemented condition. CPS I is a prominent antigen and virulence determinant that allows for survival in host blood and protection from opsonization and is required for acute virulence in *B. pseudomallei* (73). Additionally, the CPS I antigen is a current frontrunner for melioidosis diagnostic development (74), along with the previously mentioned antigen Hcp-1 (75). Thus, the nitrosative stress response in *B. pseudomallei* involves activation of some classical virulence determinants for successful eukaryotic infection while at the same time a reduction of T3SS-3 *bsa* cluster expression under the conditions and time points evaluated.

### Antibiotic resistance-associated genes

A hallmark of *Burkholderia* spp. infections is multidrug resistance via intrinsic mechanisms in these adaptable organisms. In *B. pseudomallei*, resistance mechanisms are chromosomally encoded and provide antibiotic protection via physical exclusion, enzymatic inactivation, target mutation, and importantly, many RND efflux pumps (76). Our transcriptomic analysis revealed significant differential regulation of several antibiotic resistance-associated loci in *B. pseudomallei* 1026b in the conditions tested. A putative RND efflux system spanning Bp1026b_II2076 – II2080 was significantly upregulated in its entirety, with a mean fold change ratio of 3.0 in the nitrate treatment and 2.5 in the nitrite treatment groups. However, this cluster has yet to be characterized in *B. pseudomallei* as a classical RND efflux relevant to clinical antibiotics (77), thus this expression may not be associated to increased drug resistance. Interestingly, this putative RND efflux cluster is encoded adjacent to an ABC-type export system spanning Bp1026b_II2069 – II2072 that is similarly upregulated in its entirety, 3.1-fold in the nitrate condition and 2.2-fold in the nitrite treatment group. These two export clusters are separated by a universal stress protein locus (Bp1026b_II2074) and a TetR transcriptional regulator (Bp1026b_II2075), which are similarly expressed and suggest that activation of the export systems may be reflective of a general stress response (78, 79).

Several gene loci that are directly correlated to antimicrobial resistance via enzymatic and molecular mechanisms were also identified in our analysis. Most notably, the class D β-lactamase, OXA-57 (80) encoded by Bp1026b_II2145 (*oxa*), was significantly expressed at a ratio of 4.3-fold and 3.8-fold for the nitrate and nitrite conditions, respectively. Transcription of class D β-lactamases such as OXA-57 has been shown to be increased in *B. pseudomallei* that are resistant to ceftazidime, a frontline antibiotic (81). Strikingly, the operon Bp1026b_II2141 – Bp1026b_II2145, which has been shown to be transcriptionally activated *in vivo* in response to ceftazidime treatment (82) was significantly upregulated as an entire unit (except for the response regulator *irlR2*) with an average fold change ratio of 4.2 in the nitrate condition. This operon encodes a putative regulator of the stress response (Bp1026b_II2144) and a response regulator implicated in imipenem resistance (Bp1026b_II2142, *irlR2*) in addition to the *oxa* β-lactamase amid genes for hypothetical proteins (82). Our transcriptional analyses revealed Bp1026b_II2144 to be upregulated 6.2-fold and 5.8-fold in the nitrate and nitrite conditions, respectively. Following this trend, another locus encoding a hypothetical protein of unknown function, Bp1026b_II2146, adjacent to *oxa*, was significantly upregulated 12.5-fold and 7.4-fold in the nitrate and nitrite conditions, respectively.

Additionally, the locus encoding DNA gyrase subunit A, *gyrA* (Bp1026b_I0792) was significantly downregulated −2-fold in both conditions, indicating a restriction of active cell division under the conditions tested. Furthermore, the locus encoding for dihydrofolate reductase (83), *folA* (Bp1026b_I0834) is also downregulated −2.7-fold and −2.2-fold in the nitrate and nitrite conditions, respectively, although the latter did not pass our stringent statistical threshold. In addition to the RND efflux, periplasmic β-lactamases, and LPS, among other mechanisms, *Burkholderia* spp. have been shown to acquire resistance to fluoroquinolones and trimethroprim via targeted mutations in *gyrA* and *folA*, respectively (84). Thus, our observations of significant differential regulation of key antibiotic resistance-associated gene loci implicate the nitrosative stress response in regulation of these resistance mechanisms in *B. pseudomallei*.

### Polysaccharide biosynthetic genes

In conjunction with the biofilm inhibitory phenotype resulting from nitrate and nitrite dose responses in wild type (Fig 2A, (5)), our analysis revealed significant downregulation of several gene clusters associated with polysaccharide biosynthesis. The matrix of extracellular polymeric substances (EPS) in *Burkholderia* spp. Encompasses numerous polysaccharides, adhesins, lipids, and extracellular DNA that contribute to aggregation and biofilm formation (74). To date, four capsular polysaccharide (CPS) gene clusters have been characterized in *B. pseudomallei* (74); CPS I: Bp1026b_I0499 – Bp1026b_I0524, CPS II: Bp1026b_II0468 – Bp1026b_II0480, CPS III: Bp1026b_II1956 – Bp1026b_II1966, and CPS IV: Bp1026b_I0525 – Bp1026b_I0543. Exopolysaccharide clusters homologous to *bce-I* (Bp1026b_II1956 – Bp1026b_I1966, also annotated as CPS III) and *bce-II* (Bp1026b_II1795 – Bp1026b_II1807) cepacian biosynthesis in the *Burkholderia cepacia* complex (Bcc) (85) have also been described in *B. pseudomallei* (74). Additionally, we have recently described a novel biofilm-associated biosynthesis gene cluster, *becA-R* (Bp1026b_I2097 – Bp1026b_I2927) (34). Our analysis revealed significant downregulation of CPS III/*bce-I, bce-II*, and *becA-R* biofilm-associated gene clusters, with varying degrees of cluster coverage, among the differentially expressed datasets from both the nitrate and nitrite conditions.

The CPS III cluster displayed a reduced expression of −2.2 mean fold change for six transcripts in the nitrate comparison, and −2.7 mean fold change for seven transcripts in the nitrite comparison. Similarly, the *bce-II* cluster displayed reduced expression of −2.5 mean fold change for seven transcripts in the nitrate comparison, and −2.9 mean fold change for six transcripts in the nitrite comparison. Within the *becA-R* biofilm cluster, Bp1026b_I2922 (*becM*) was downregulated −5.2-fold and Bp1026b_I2923 (*becN*) −7.3-fold in the nitrate condition, with *becN* also down −7.1-fold in the nitrite condition. Lastly, Bp1026b_II0477, encoding a predicted ADP-heptose-LPS-heptosyltransferase in CPSII was downregulated −2.3-fold in the nitrate condition. Altogether, these results demonstrate the transcript level reduction of key EPS matrix components, notably exopolysaccharide clusters, that go hand in hand with the biofilm inhibition response to both nitrate and nitrite supplementation in *B. pseudomallei*. The reduction of expression in these biofilm-associated clusters is also dependent on the intact NarX-NarL system, as all the above trends are reversed in pairwise comparisons of Δ*narX* and Δ*narL* to the wild type under nitrate treatment.

### Secondary metabolite biosynthetic gene clusters

In addition to polysaccharide biosynthetic clusters, *Burkholderia* spp. encode several gene clusters that produce antimicrobial natural products (86). *B. pseudomallei* 1026b encodes a combination of 15 nonribosomal peptide synthetase (NRPS) and polyketide synthase (PKS) biosynthetic gene clusters residing on both chromosomes (61). Our analysis identified four clusters on chromosome II in which all loci were similarly differentially regulated in both the nitrate and nitrite stress comparisons (Fig 7). Bactobolin is an antibiotic encoded by a 120-Kb DNA element in *B. pseudomallei* and *B. thailandensis* that is responsive to AHL mediated quorum sensing (87). The bactobolin biosynthetic cluster, encoded by loci Bp1026b_II1232 – Bp1026b_II1251 (*btaA – btaU*) (88), was markedly upregulated at a mean fold change of 3.1 in the nitrate treatment group and 3.0 in the nitrite treatment group. In both comparisons, all gene loci in the bactobolin cluster were among the most significantly expressed with consistently low false discovery rates.

**Fig 7.**
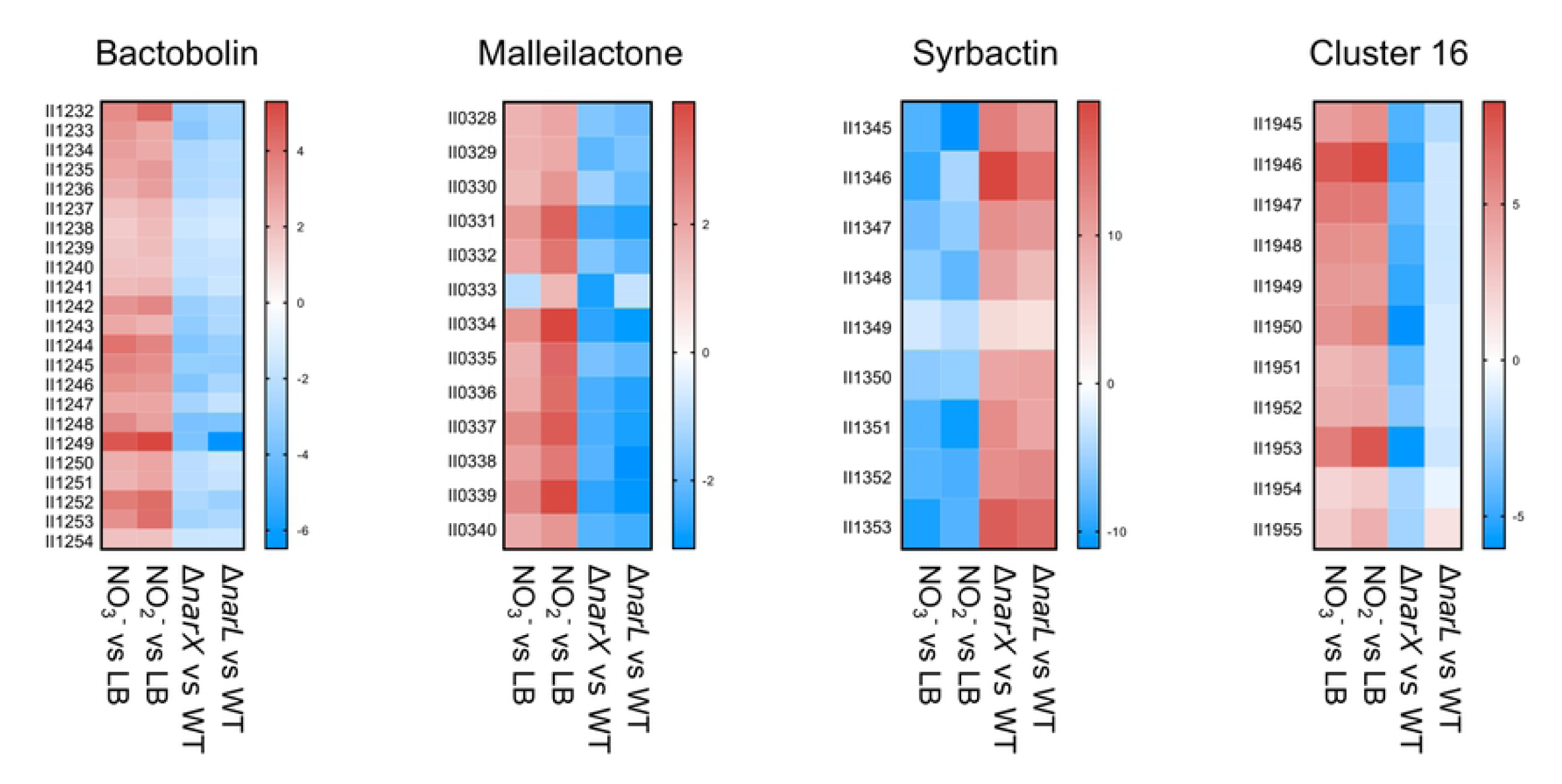
Key clusters associated with antibiotic production, cytotoxic siderophore synthesis, and proteasome inhibition are differentially regulated by nitrate and nitrite. Differential regulation of secondary metabolite synthesis clusters, bactobolin, malleilactone, syrbactin, and the cryptic cluster 16. Data was extracted from DESeq2 analysis of significantly regulated gene loci and pooled as mean fold change values, which are visualized here. Uniform regulation is evident among the treatment conditions, with nitrate and nitrite conditions similarly regulating clusters opposed by Δ*narX* and Δ*narL* mutant trends in the nitrate treatment condition. Color density gradients represent up-regulation (red), down-regulation (blue), and no differential regulation (white).

Another biosynthetic gene cluster that was identified as significantly upregulated in both nitrate and nitrite stress conditions encodes for malleilactone, a cytotoxic siderophore (89). The malleilactone biosynthetic cluster contains 13 ORFs (Bp1026b_II0330 – Bp1026b_II0341), including two large polyketide synthase gene loci (*malA* and *malF*) and a LuxR-type transcription factor (*malR*), indicating that its expression is mediated by an AHL quorum-sensing system (89). When comparing wild type in the nitrate-supplemented media versus regular media, 7/13 of the malleilactone genes were significantly upregulated by a mean fold change ratio of 2.4, and in the nitrite condition versus regular media, 11/13 loci are significantly expressed at a fold change ratio of 3.1. Interestingly, malleilactone is among the few PKS/NRPS clusters that is conserved among the Bpc, including *B. mallei* and *B. thailandensis*, the latter of which has been shown to be important for virulence (61, 89). Thus, in apparent coordination with bactobolin antibiotic expression, *B. pseudomallei* similarly regulates the transcription of the malleilactone siderophore biosynthetic genes as a response to the biofilm-inhibitory doses of nitrate and nitrite.

In stark contrast to bactobolin and malleilactone upregulation in these conditions, a cluster of genes from Bp1026b_II1345 – Bp1026b_II1353 (*syrA – syrI*) encoding syrbactin (61), was significantly downregulated in both the nitrate and nitrite treatment groups. Syrbactin is a proteasome inhibitor produced by a hybrid PKS/NRPS cluster comprised of nine genes. All nine genes in the syrbactin cluster were downregulated at a mean fold change of −7.1 in the nitrate treatment group and −7.5 in the nitrite treatment group. For both treatment groups, syrbactin biosynthetic cluster genes were among the most downregulated genes in the differential expression analysis. However, analysis of Δ*narX* and Δ*narL* under nitrate stress compared to the wildtype reversed this trend, implicating the NarX-NarL system in regulation of secondary metabolism in *B. pseudomallei* 1026b. Further supporting this hypothesis, a homolog of the recently described global regulator of secondary metabolism (90), Bp1026b_I0582 (*scmR*), is significantly downregulated −2.0-fold in the nitrate treatment group and −1.7-fold in the nitrite group. ScmR is a LysR-type transcriptional regulator that represses many biosynthetic gene clusters in the Bpc (90). In both analyses of Δ*narX* and Δ*narL* under nitrate stress, *scmR* is notably upregulated, 2.6-fold and 2.9-fold, respectively. Collectively, these results suggest that nitrosative stress induces transcription of bactobolin antibiotic and malleilactone siderophore biosynthesis and represses transcription of the syrbactin proteasome inhibitor via the global regulator *scmR*, which is dependent on a functioning NarX-NarL two-component system.

### Genes associated with the stringent response

The growth conditions tested in these experiments causing nitrosative stress in sessile cells resulted in the activation of several genes associated with the stringent response. In β- and γ-proteobacteria, enzymes RelA and SpoT regulate production of (p)ppGpp, a messenger molecule that signals nutritional stress and regulates transcription and cell physiology (91), including biofilm formation (92). Importantly, *B. pseudomallei* encodes homologs of both these ppGpp-metabolic loci (*relA*, Bp1026b_I1913; *spoT*, Bp1026b_I0746), the deletion of which have been considered for vaccine candidacy due to attenuation in both acute and chronic melioidosis models (93). Our results comparing sessile *B. pseudomallei* cells subject to elevated nitrate and nitrite levels revealed significant upregulation of *relA* as well as several loci associated with oxidative stress, destabilization of rRNA/tRNAs, reduction of DNA replication, and general stasis survival genes (S2 Fig). The *relA* locus, encoding the GTP hydrolase associated with (p)ppGpp alarmone production (94), was significantly upregulated at a fold change ratio of 2.1 in both the nitrate and nitrite treatment conditions. The relative expression of *relA*, along with several other key transcripts identified in this study (Fig 8), matched observed trends from the larger DESeq2 analysis.

**Fig 8.**
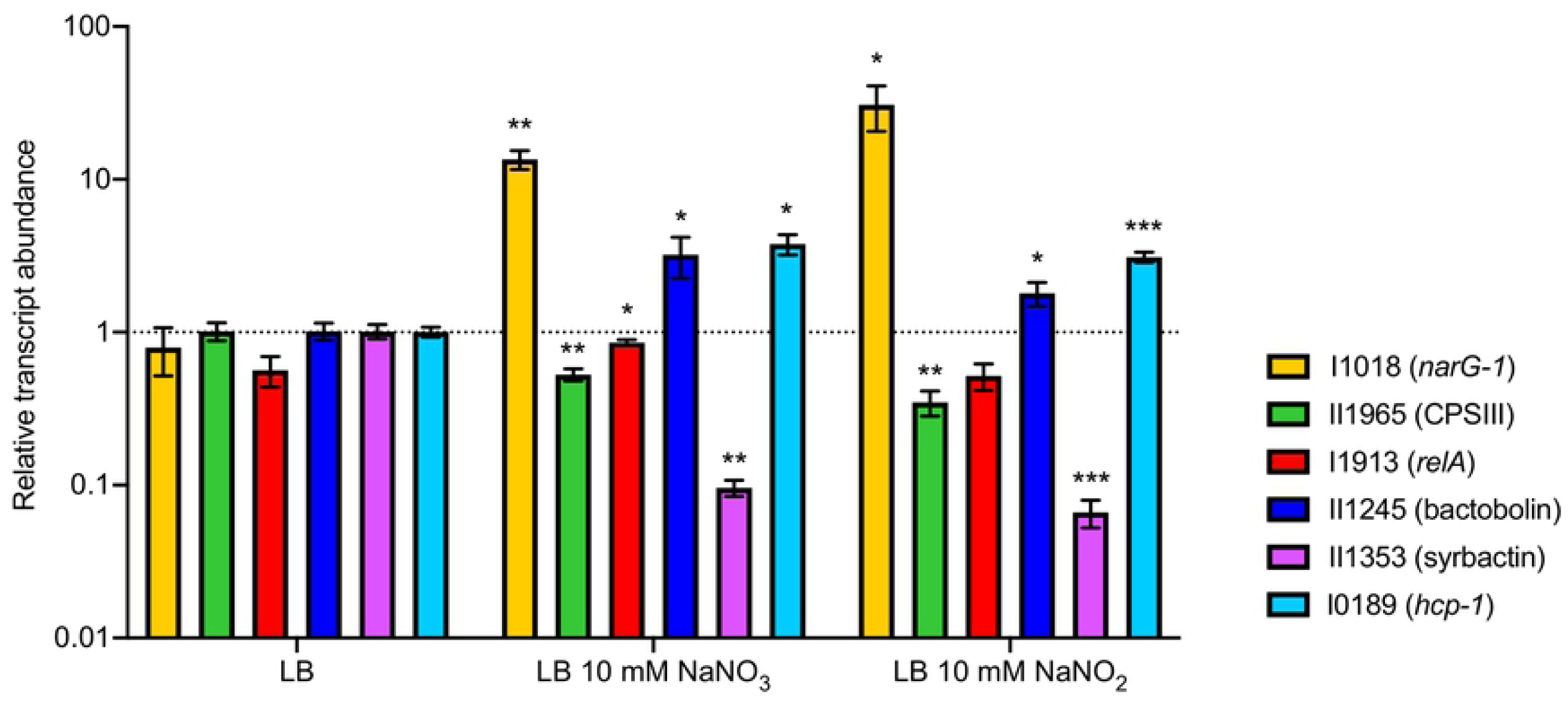
Quantitative evaluation of key transcripts confirms trends in the DESeq2 data set. Relative abundance of transcripts differentially regulated by both nitrate and nitrite treatment conditions as compared to baseline expression in LB. Fold change in transcript level was calculated using the Pfaffl method and normalized to the housekeeping transcript for 23S rRNA. Statistical significance was determined using a one-tailed heteroscedastic Student’s T-test (* = p<0.05, ** = p<0.01, *** = p<0.001).

Several rRNA loci encoding 50S and 30S subunits were significantly downregulated in both conditions and tRNA encoding-loci were among the most significantly downregulated in the nitrate treatment group. The downregulation of rRNA/tRNA loci in response to nitrate and nitrite treatment corresponds to potential markers of *in vivo* infections as described in a recent transcriptomic profile of *B. pseudomallei* (82). Similarly, the locus encoding for the translation initiation factor IF1, Bp1026b_I2473, was significantly downregulated −7.0-fold in the nitrate condition and −4.5-fold in the nitrite condition. Consistent with the observed reduction in expression of loci associated with translation, ribosomal subunits, and transcription, the downregulation of *gyrA* further supports the link between nitrate and nitrite stress with the stringent response. Collectively, these data indicate a general downregulation of genes associated with bacterial replication and protein synthesis, reminiscent of the stringent response in *E. coli* (95).

### *B. pseudomallei* nitrate sensing mutants are deficient in intracellular survival in mouse macrophages

Successful intracellular pathogens require the ability to survive within phagocytic cells such as macrophage that bombard intruders with reactive oxygen and nitrogen intermediates (21, 29). To determine the ability of the nitrate sensing-deficient mutants to infect and survive in a hostile macrophage environment amid nitrosative stress, a murine macrophage cell line (RAW 246.7) was co-incubated with wild-type, Δ*narX*, and Δ*narL* strains at a MOI of ∼2. The number of intracellular bacteria was determined after 2h, 6h, and 20h post-exposure when eukaryotic cells were lysed and plated to count bacterial colony forming units (CFU). At 2h, there was a notable increase in bacterial internalization into the murine macrophage cells; however, at 6h and 20h Δ*narX* and Δ*narL* mutants were significantly impaired at intracellular survival (Fig 9A). Two hours after infection, the wild type was recovered at lower titers than either of the two mutant strains (Fig 9B). Δ*narX* and Δ*narL* were internalized at 166% and 168% compared to the wild type, respectively, suggesting possible preferential uptake of these mutant bacteria by the murine macrophage. At 6h post-infection, Δ*narX* colony forming units were recovered at 56%, and Δ*narL* at 33% compared to the wild type, indicating an early-onset defect in intracellular replication. After 20h of infection, the observed Δ*narX* and Δ*narL* defects were maintained and CFU were recovered at 51% and 33% compared to wild type, respectively (Fig 9A). When comparing Δ*narX* and Δ*narL* mutants to wild type infection dynamics, CFU values were statistically significant at all three time-points of infection, however Δ*narL* was more drastically attenuated in this model. Thus, these results indicate that although attachment and internalization of the mutant strains tested are elevated at 2h post-infection, there are significant defects in *B. pseudomallei* lacking the NarX sensor kinase and the NarL DNA-binding regulator during intracellular replication and long-term survival.

**Fig 9.**
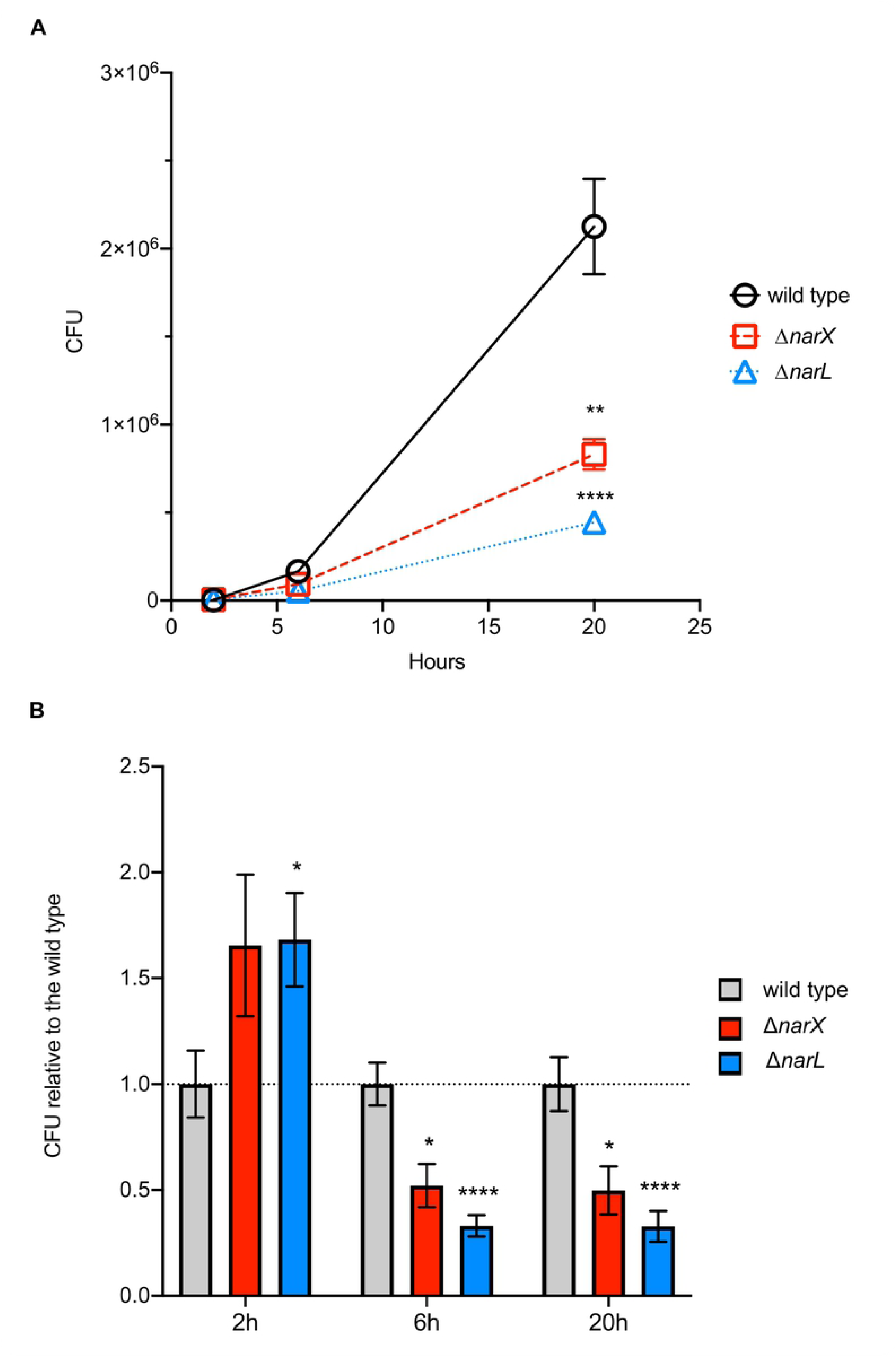
*B. pseudomallei* mutants lacking NarX and NarL are deficient in intracellular replication despite being internalized at a higher rate than the wild type. RAW264.7 cell monolayers were infected at an MOI of ∼2 with three strains (wild type, Δ *narX*, and Δ *narL*) and intracellular survival was measured at 2h, 6h, and 20h post-infection. At 1h post-infection, cells were treated with 750 μg/mL kanamycin to kill extracellular bacteria. Total CFU levels were calculated (A) as well as CFU relative to the wild type (B) at all time-points. The internalization efficiency of the mutant is evident at 2h post-infection, yet the intracellular replication efficiency is hindered at 6h and infection is attenuated at 20h (B). Wild type is depicted in black (A) or grey (B), with Δ *narX* in red and Δ *narL* in blue throughout. Statistical significance was determined using the Holm-Sidak method across multiple Student’s T-tests (* = p<0.05, ** = p<0.01, **** = p<0.001).

## Discussion

Understanding the biofilm dynamics of *B. pseudomallei* in the context of environmental sensing of nitrate and nitrite has broad implications for transmission epidemiology as well as the pathogenicity and clinical mitigation of this facultative intracellular organism. The metabolic versatility and adaptability of *B. pseudomallei* underscores its success as a ubiquitous tropical saprophyte (22) as well as a human pathogen capable of infecting all tissue types tested (96). *B. pseudomallei* is recognized as a potential threat to people who come in contact with contaminated soil from domestic gardens in endemic regions (97, 98), and may be linked to anthropogenic sources of nitrate and urea in these gardens (99, 100). The present study is based on characterization of the nitrosative stress response in *B. pseudomallei* as it relates to biofilm inhibition via the nitrate-sensing NarX-NarL two-component regulatory system. Using comparative transcriptome analyses between conditions of nitrate and nitrite treatments, we identified the global regulatory mechanisms of nitrate sensing that are intimately linked to both physiology and pathogenicity in *B. pseudomallei*. Our previous study identified a link between genes in the *narX-narL-narGHJI*_*1*_*-narK*_*1*_*-narK*_*2*_ regulon, biofilm inhibition and reduction of cyclic di-GMP levels (5). We further characterized the *narX-narL* system in conjunction with biofilm inhibition and discovered a complex regulatory system encompassing key elements of the stringent response, secondary metabolism, virulence, and antibiotic tolerance (Fig 10).

**Fig 10.**
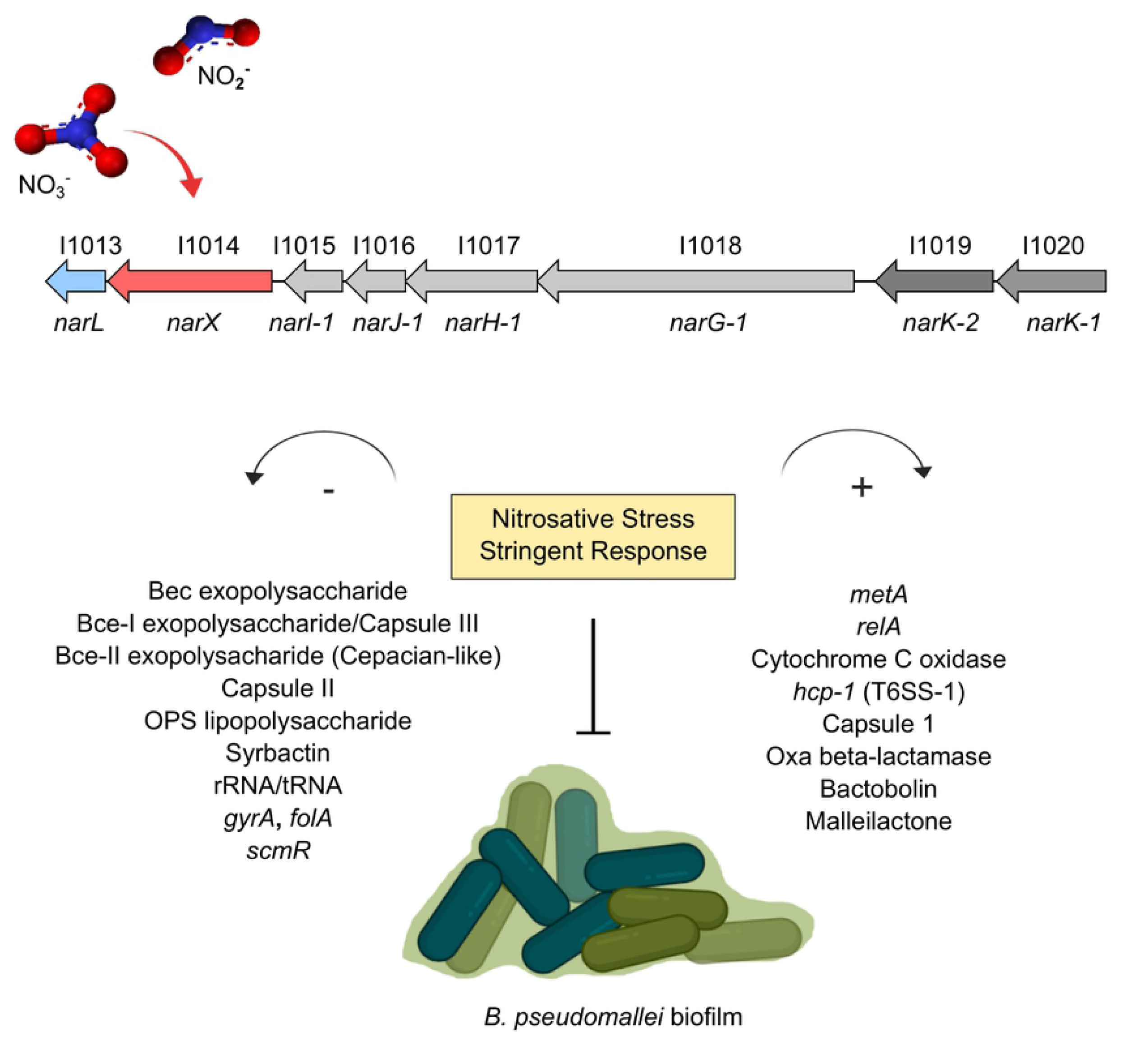
Working model of nitrate-dependent biofilm inhibition and activation of the stringent response in *B. pseudomallei*. Nitrate, and to a lesser extent nitrite, activate the *narXL* sensing mechanism in *B. pseudomallei* which leads to metabolism of N-oxides, activation of nitrosative stress and stringent responses, and indirectly inhibiting biofilm formation. Several biofilm-associated gene clusters such as capsules, exopolysaccharides, and lipopolysaccharides are down-regulated along with several housekeeping genes that are necessary for growth and division. On the other hand, secondary metabolic biosynthesis clusters are up-regulated in conjunction with pathogenicity-associated genes, an alternative respiration mechanism, as well as the classical indicator for stringent response, *relA*, and methionine metabolism gene *metE*.

Our initial observation of a biofilm inhibition phenotype that is comparable in both nitrate and nitrite treatment in oxic conditions is complicated by the fact that nitrite-dependent biofilm inhibition does not require the NarX-NarL system (Fig 2A). This disparity lead us to hypothesize that the NarX-NarL system in *B. pseudomallei* can discriminate between nitrate and nitrite ligands, with clear preference towards nitrate, as evidenced by NarX protein characterization studies in *E. coli* (9). This apparent ligand preference is further demonstrated in our transcriptomic analyses including Δ*narX* and Δ*narL* mutants in the presence of either nitrate (Figs S3B and S3E) or nitrite (Figs S3C and S3F). In response to nitrate, both mutants activate and upregulate 449 transcripts (7% of *B. pseudomallei* coding sequences), with 234 of those shared among them (Fig S3B), and downregulate 353 (6% of CDS), with 184 overlapping in regulation, using stringent significance thresholds. Conversely, Δ*narX* and Δ*narL* mutants compared to the wild type in the presence of nitrite only differentially regulated 10 transcripts in either a positive or a negative manner, further suggesting that exogenous nitrite is not regulated by NarX-NarL (Fig S3C, S3F). Future studies are required to characterize the complex regulatory systems that facilitate both nitrate and nitrite signals in *B. pseudomallei*, although this is beyond the scope of the current study. Sensing of similar N-oxides, nitrate and nitrite, are facilitated by twin two-component systems NarX-NarL and NarQ-NarP in enteric bacteria (101), of which *B. pseudomallei* only encodes NarX-NarL (9). Given that our previous *in silico* analyses did not discover a duplication of the NarX-NarL system on chromosome II in *B. pseudomallei* (5), and given the data presented here, we propose that the NarX-NarL system preferentially binds nitrate as a ligand.

The NarX-NarL system regulates biofilm formation in a nitrate-dependent manner, through direct or indirect downregulation of key EPS matrix components. Our analyses also implicate the stringent response, as activated by the nitrosative stress conditions tested, in regulation of the biofilm cycle. Recently, the stringent response mediated by the ppGpp synthases RelA and SpoT, were implicated in a nutrient-starvation model of biofilm dispersal described in *Pseudomonas putida* (102). The stringent response has been linked to nitrogen starvation in *E. coli* via σ^54^-activated *relA* transcription (103), whereas our results describe upregulation of *relA* in response to nitrosative stress. Additionally, several genes regulated by the alternative sigma factor σ^54^ such as the virulence-associated *bkdA1-bkdA2-bkdB-lpdV* operon (104) are significantly upregulated in our pairwise comparisons of both nitrate and nitrite treatment groups to the wild type. The *bkdA1-bkdA2-bkdB-lpdV* cluster encodes a keto acid dehydrogenase that has been identified in transcriptomic studies of planktonic *P. aeruginosa* compared to developing biofilms (105, 106). These differential expression patterns are not surprising considering σ^54^ is responsible for nitrogen regulation and is responsive to ppGpp signaling in *E. coli* (95). In contrast to the classical nutrient starvation model of stringent response in antibiotic tolerant biofilms (107), we propose a similar outcome due to nitrosative stress in *B. pseudomallei*. Indeed, the responses to both nitrate and nitrite stress in conjunction with biofilm inhibition, revealed coordinated expression of virulence-associated loci, repression of nonessential growth and activation of antibiotic tolerance factors. However, it is worth taking these transcriptional trends with a grain of salt, considering that expression of genes required for physiological factors such as efflux pumps can be induced during generalized stress responses (108).

Nonetheless, the *in vitro* transcriptional response to nitrosative stress provides the foundation to understand physiological response in the context of both environmental and *in vivo* infection scenarios. Facultative intracellular pathogens must withstand several innate immune response molecules, such as reactive oxygen and nitrogen species, sparking an interest in potential intrinsic genetic and enzymatic resistance mechanisms in bacteria. Resistance to RNI has been shown to involve methionine and homocysteine metabolism in *M. tuberculosis* and *Salmonella typhimurium* (30, 60, 109), which couples with our observation that *metE* is the most significantly upregulated locus in both nitrate-supplemented and nitrite-supplemented conditions. Further mutational analyses will be necessary to determine if transcripts identified in this study are implicated in response to RNI. To bridge this gap, we analyzed the fitness of Δ*narX* and Δ*narL B. pseudomallei* 1026b mutants in eukaryotic cell infection, and observed a significant defect in intracellular survival efficiency in both mutants (Fig 9). Although susceptible to the bactericidal activity of IFN-γ-stimulated macrophages, with RNI having a stronger effect than ROI (29), *B. pseudomallei* can survive intracellularly in phagocytic cells (110). Our observation of replication-deficient *B. pseudomallei* lacking either the NarX histidine kinase or the NarL DNA-binding regulator suggests that nitrate metabolism is important for intracellular survival. The intracellular growth deficiencies of Δ*narX* and Δ*narL B. pseudomallei* is best explained by lack of active NarG and subsequent nitrate reductase activity, which is important for *Mycobacterium tuberculosis* intracellular growth (57), although this definitive connection remains to be characterized in *B. pseudomallei*. Intriguingly, conversion of nitrate to nitrite in *M. tuberculosis* leads to growth retardation (52), which is similar to the trend we observed in *B. pseudomallei*.

The differing responses to nitrate and nitrite in our *B. pseudomallei* anaerobic growth model (Fig 2C) raises the possibility of a disparity between exogenous nitrite and endogenously-produced nitrite via nitrate respiration, as seen in *M. tuberculosis* (52). Opposing this theory; however, our transcriptomic analyses of exogenous nitrate and nitrite revealed a strikingly similar genome-wide response. One possible explanation for this observation is that our global analysis is based on a transcriptional response from cells grown in oxic conditions, although oxygen limitation is a hallmark of biofilms that shift towards anaerobic metabolism during infections (111). Nonetheless, the responses of *in vitro* biofilms to nitrate and nitrite were strikingly similar in our study (Fig 5A, 5B). The transcriptional trends identified by the nitrate stress response are dependent on both components of the NarX-NarL system, as the differential expression patterns of both Δ*narX* and Δ*narL* in the nitrate condition were opposite to that of the wild type (Fig 7).

Among the clearest examples of a similar response to both N-oxides is the regulation of secondary metabolism biosynthetic gene clusters (BGCs). Transcriptional activation of the complete bactobolin and malleilactone clusters as well as the cryptic cluster 16, coupled with the downregulation of the complete syrbactin biosynthesis cluster follow the same trends for either nitrate and nitrite, yet are dependent on a functioning NarX-NarL system (Fig 7). In *B. thailandensis*, a global repressor of BGCs, MftR regulates both bactobolin and malleilactone production as well as *narX* and *narL*, and the T3SS regulator *bsaN*, suggesting that BGC regulation is inherently linked to host environment adaptation and ultimately virulence (112). In *B. pseudomallei* bactobolin and malleilactone are quorum sensing (QS)-controlled secondary metabolites, which may have implications for the intricate QS regulation of virulence in this organism (62). Our results potentially link these key QS-regulated factors to the stringent response as activated by nitrosative stress in *B. pseudomallei*. The possibility that the stringent response is linked to quorum sensing in *B. pseudomallei* is not surprising, considering the strong association between these two systems in *P. aeruginosa* (113, 114). In this context, we hypothesize that nitrosative stress activates QS-regulated metabolites that can aid *B. pseudomallei* in adapting to host environments.

The hypothesis that *B. pseudomallei* can maintain infection during anaerobic conditions through nitrate sensing and denitrification is in part supported by a recent genome-wide analysis of environmental and clinical strains of *B. cenocepacia*, a phylogenetic relative (115). Wallner et al. noted that out of 303 *B. cenocepacia* strains analyzed, the *narX-narL* nitrate sensing system is present in all clinically-derived strains and the *narGHJI* operon is variably present among clinical and environmental strains (115). Our characterization of intracellular replication deficiencies in both *narX* and *narL* mutants of *B. pseudomallei* in macrophages warrants further investigation and development of inhibitory mechanisms for this two-component system. Indeed, such two-component systems have been recognized as promising targets for antimicrobial drug development, given their importance for bacterial survival and pathogenicity in the host (116). Several two-component systems have been described to be involved in virulence in pathogenic *Burkholderia* spp. including *B. pseudomallei*, reinforcing the need to characterize these signaling and gene-regulation pathways to assess their role in *Burkholderia* host-associated ecology (117). To this end, we describe the transcriptome of a clinical isolate, *B. pseudomallei* 1026b, during *in vitro* nitrate- and nitrite-supplemented growth conditions and characterize a signal transduction network reliant on a functional *narX-narL* two-component system. Furthermore, our investigation of this system in an *in vivo* infection model highlights its importance as a virulence-facilitating factor in *B. pseudomallei*. Future work should continue investigations into the *narX-narL* system during infection of more complex model systems, as well as the regulation of secondary metabolite biosynthetic clusters in nitrate-rich environments. Targeting the *narX-narL* system in a host-associated context will provide valuable information for mitigating *B. pseudomallei* infection and disease progression.

## Acknowledgements

We thank Grace Borlee and Ian McMillan for critical review and discussions of this manuscript. We are also grateful to Justin Lee, Marylee Layton, Mark Stenglein, and the MIP NGS Illumina Core for technical assistance with library prep and sequence generation.

---

**Fig S1. Genomic conservation of the Nar regulon in the Bpc**. *Burkholderia pseudomallei* 1026b, *B. mallei* ATCC23344, and *B. thailandensis* E264 nitrate reduction regulons are compared here. Orthologous sequences were extracted and aligned using EasyFig (118) in Python v2.7. Illustrations depict results of blastn annotations represented by colored bars spanning chromosomal segments, with minimum percent identity of 0.60 and a threshold E-value of 1E-3. Yellow-to-red bars depict sequence homology amid inverted sequences on a color density gradient indicating percent homology, whereby dark red indicates the most homology.

**Fig S2. Overview of the transcriptomic analysis of nitrate/nitrite-mediated biofilm inhibition of *B. pseudomallei***. 36 individual cDNA libraries were generated from total RNA (Sample Processing), before single-end sequencing (Illumina Sequencing) and statistical analysis (Bioinformatics). 409 million reads were generated, from which statistically significant differential expression analysis was derived. Following quality assessment of reads, mapping and alignment to the *B. pseudomallei* 1026b genome, DESeq2 statistical analysis was performed in R.

**Fig S3. Nitrate and nitrite treatments differentially regulate similar transcripts in wild-type *B. pseudomallei*, while *narX* and *narL* mutants respond to nitrate but not nitrite treatments**. (A, D) Wild-type *B. pseudomallei* in either the nitrate (NO_3_^-^) or nitrite (NO_2_^-^) treatment condition. Upregulated transcripts: nitrate vs. LB = 274, nitrite vs LB = 237; (D) downregulated transcripts: nitrate vs. LB = 316, nitrite vs LB = 239. (B, E) Δ*narX* and Δ*narL* strains in the nitrate (NO3 ^-^) treatment condition. (B) Upregulated transcripts: Δ*narX* vs. wild type = 349, Δ*narL* vs. wild type = 334; (E) downregulated transcripts: Δ*narX* vs. wild type = 285, Δ*narL* vs. wild type = 252. (C, F) Δ*narX* and Δ*narL* strains in the nitrite (NO2 ^-^) treatment condition; (C) upregulated transcripts: Δ*narX* vs. wild type = 1, Δ*narL* vs. wild type = 0; (F) downregulated transcripts: Δ*narX* vs. wild type = 8, Δ*narL* vs. wild type = 4.

**Fig S4. Functional characterization of transcriptionally active and differentially regulated genes reveals defects in translation, ribosomal structure and biogenesis**. Cluster of orthologous groups comparisons of up- and down-regulated loci from both nitrate (A) and nitrite (B) treatment conditions.

**Fig S5. Clusters of orthologous groups assigned to the significant differentially regulated transcripts identified in this study**. Functional characterization of the differentially expressed genes in response to nitrate-supplemented and nitrite-supplemented treatment.

## Funding

MRM was supported by an NSF NRT-GAUSSI fellowship; Results presented in this paper are based upon collaborative work supported by a National Science Foundation NRT Grant No. 1450032. Any opinions, findings, conclusions, and recommendations expressed in this paper are those of the author(s) and do not necessarily reflect the views of the National Science Foundation.

## Author Contributions

Conceptualization and experimental design: MRM and BRB. Performed the experiments: MRM. Analyzed the data: MRM. Project administration, resources, supervision, and validation: BRB. Wrote and edited the paper: MRM and BRB.

## Data Availability

Raw sequence files associated with this study are deposited at the European Nucleotide Archive (http://www.ebi.ac.uk/ena) under the primary accession number PRJEB38907.

